# The addiction-susceptibility TaqIA/Ankyrin repeat and kinase domain containing 1 kinase (ANKK1) controls reward and metabolism through dopamine receptor type 2 (DR2)-expressing neurons

**DOI:** 10.1101/2022.08.12.503577

**Authors:** Enrica Montalban, Roman Walle, Julien Castel, Anthony Ansoult, Rim Hassouna, Ewout Foppen, Xi Fang, Zach Hutelin, Sophie Mickus, Emily Perszyk, Anna Petitbon, Jérémy Berthelet, Fernando Rodrigues-Lima, Alberto Cebrian-Serrano, Giuseppe Gangarossa, Claire Martin, Pierre Trifilieff, Clémentine Bosch-Bouju, Dana. M Small, Serge Luquet

## Abstract

Significant evidence highlights the importance of genetic variants in the development of psychiatric and metabolic conditions. Among these, the Taq1A polymorphism is one of the most commonly studied in psychiatry. TaqIA is located in the gene that codes for the Ankyrin repeat and kinase domain containing 1 kinase (ANKK1) near the dopamine D2 dopamine receptor (DR2) gene. Depending on race it affects 30 to 80% of the population and its homozygous expression of the A1 allele correlates with a 30 to 40% reduction of striatal DR2, a typical feature of addiction, over-eating and other psychiatric pathologies. The mechanisms by which the variant influences dopamine signaling and behavior is unknown. Here we used transgenic and viral-mediated strategies to reveal the role of ANKK1 in the regulation of activity and functions of the striatum. We found that *Ankk1* is preferentially enriched in striatal DR2 expressing neurons and that Ankk1 loss-of-function in dorsal and ventral striatum leads to alteration in learning, impulsive, and flexible behaviors resembling the endophenotypes described in A1 carriers. We also observed an unsuspected role of ANKK1 in striatal DR2-expressing neurons in the ventral striatum in the regulation of energy homeostasis and documented differential nutrient partitioning in humans with versus without the A1 allele. Overall, our data demonstrate that the Ankk1 gene is necessary for the integrity of striatal functions and reveal a new role for ANKK1 in the regulation of body metabolism.

## Introduction

Psychiatric conditions are multifactorial diseases caused by both genetic and environmental factors. Even though classically considered as distinct diseases, various psychiatric disorders share common symptomatic dimensions such as alterations of mood, cognitive functions or reward processing, suggesting similar pathophysiological mechanisms. In line with this, the Research Domain Criteria (RDoC) proposes to classify psychiatric illnesses based on core neurobiological, behavior, genetic dimensions, aiming at identifying both the mechanisms that are shared across multiple psychiatric disorders, as well as the processes that are unique to specific psychiatric symptoms^1,2^. Interestingly, there is increasing evidence that patients with psychiatric disorders also have disturbances in energy metabolism and a higher risk of developing metabolic syndrome, with appetite changes as core feature of multiple diseases^3,4^. This raises the intriguing idea of the existence of an overlap in the pathogenic mechanisms that underlie neuropsychiatric and metabolic symptoms.

Over the past 20 years, Genome Wide Association Studies (GWAS) allowed the identification of many genetic alterations underpinning psychiatric disorders^5^. This includes single nucleotide polymorphism (SNPs), the most common type of existing genetic variation. Among SNPs, TaqIA polymorphisms have attracted growing attention. TaqIA is located in the gene that codes for the Ankyrin repeat, and kinase domain containing 1 kinase (ANKK1) and corresponds to the single nucleotide polymorphism A2 (T→C) in the position 2137 of *Ankk1* transcript. The TaqIA variant results in an amino acid change (E[GAG]→K [AAG],Glu→Lys) in position 713 of Ankk1 protein in humans. The minor A1 variant is the ancestral polymorphism while the A2 variant only recently appeared in primate evolution^6^. ANKK1 maps on the NTAD (NCAM1-TTC12-ANKK1-DR2) gene cluster, which comprises the DA receptor D2 (DR2), the tetratricopeptide repeat domain 12 (TTC12) and the neural cell adhesion molecule 1 (NCAM1) genes. The four genes have been associated with different psychiatric pathologies although the strongest association has been made with addiction and dysregulation of feeding behaviors. TaqIA corresponds to three variants, A1/A1, A1/A2 and A2/A2. Approximately 30% of European, up to 80% of Asian, and 40% of African populations possess one or two copies of the A1 allele. Homozygous dosage of the A1 allele correlates with a 30∼40% reduction of striatal DR2 abundance^7^. Yet, such a decrease in DR2 availability is a typical feature of addiction and believed to be a core endophenotype responsible for compulsive drug consumption^8^, overeating^9,10^ and obesity-related reduction in activity^11^.

Strikingly, in humans the A1 allele not only relates to several psychiatric and neurological disorders such as attention deficit, hyperactivity disorder^12^, Parkinson’s disease^13^, and addiction^14,15,16,17,18,19^, but also to metabolic dysfunction and eating disorders^20,21,22,23^. Indeed, A1 carriers are more likely to have increased waist circumference^24^ and risk for obesity^25,20,21^. At the other side of the spectrum, recent studies have also reported an association between the presence of A1 and some of the features characteristic of anorexia nervosa^22^. Of note, weight loss has been reported to be easier in obese individuals bearing the A1 variant^23^. Altogether, this suggests that A1 might be a genetic node for the convergence of neuropsychiatric and metabolic symptoms.

In addition, alterations in reward processing, motivation, working memory and cognitive flexibility are observed across all the pathologies associated to TaqIA^26,27,28,29^. From a neurobiological point of view, such symptomatic dimensions have all been associated with a dysregulation of the cortico-limbic system, and its regulation by dopamine (DA) neurotransmission^30,31^. Accordingly, the A1 variant is associated with reduced activity in the prefrontal cortex and striatum during reversal learning^26^, reduced activity in the midbrain, prefrontal cortex and thalamus during consumption of milkshake^32^, greater impulsivity^33,34,35^ or steeper delayed discounting^36^, which are typical features of several psychiatric symptoms and eating disorders.

These observations strongly suggest that; I) the interaction between environment and Ankk1 is critical in the susceptibility to reward-related and metabolic-based dysfunctions, and that ii) perturbations of various components of ingestive behavior in A1 carriers may result from a dysregulation of DR2-dependent DA transmission. To date, the molecular and cellular functions of Ankk1 remain largely unknown primarily due to the lack of animal models. As a consequence, very little is known regarding the molecular mechanisms by which A1 and A2 variants of ANKK1 alter DA signaling and, in general, in which direction those variants affect Ankk1 activity and DR2 signaling to contribute to the protection or the vulnerability to psychiatric and metabolic diseases. Herein, we show that *Ankk1* m-RNA is highly expressed in both the nucleus accumbens (NAc) in the ventral part of the striatum, and the dorsal striatum (DS), with a selective enrichment in the striatal DR2-spiny projection neurons (SPNs). Using different mouse models, we explored the behavioral and metabolic consequences of either Ankk1 downregulation in DR2-expressing neurons, or Ankk1 loss-of-function in dorsal striatal and accumbal territories. We show that loss-of-function of Ankk1 in the NAc is sufficient to recapitulate the main features of Ankk1 downregulation in DR2-expressing neurons and also mediates metabolic changes in whole body nutrient utilization. Given that the metabolic phenotype was unsuspected and not explored in humans, we also performed a translational study and in accordance with the mouse data, found evidence for differential whole-body metabolism in A1 carriers versus noncarriers. This work provides the first reverse translational approach exploring the biological functions of Ankk1 in the central regulation of both metabolic and reward functions and further translates the metabolic phenotype discovered in mice to humans. Collectively, our data show that Ankk1 loss of function is sufficient to mimic some of the phenotypic characteristics of Taq1A individuals and reveal a new role of Ankk1 and the NAc in regulating energy metabolism.

## RESULTS

### Striatal regional distribution and regulation of Ankk1-mRNA

Consistent findings in humans reveal that TaqIA polymorphisms are associated with alterations in the activity of the brain reward system, and in particular in striatal circuits^37^. We first tested and confirmed the expression of *Ankk1* m-RNA in both the NAc and the DS and an enrichment in the DS as compared to the NAc (**Fig.1-A**^**I**^). Then, to gain insights on the cellular expression of *Ankk1* in the DR1- or the DR2-SPNs of the striatum, we re-analyzed available RNA-seq from striatal ribosome affinity purification (TRAP) technology (**Fig.1-B**)^38^ and showed that *Ankk1* m-RNA is virtually absent in DRD1-SPNs but selectively expressed in DR2-SPNs (**Fig.1-B**^**I**^). Next, we showed that stimulation of DA receptors by apomorphine treatment (**Fig.1-C)** induced sustained downregulation of *Ankk1* m-RNA in both NAc and DS 1h and 3h after injection (**Fig.1-C**^**I**^, **C**^**II**^).

The finding that in the striatum Ankk1 is selectively expressed in DR2-SNPs led us to assess whether some of the phenotypes of A1 carriers could originate from alterations of the functions of Ankk1 selectively in DR2-expressing neurons. To do so, we generated Ankk1^lox/lox^ mice in which exon 3-8 are flanked with LoxP sites allowing for CRE-targeted deletion of most of the Ankk1 gene (**Fig.1-D and Supp.1**), that we crossed with a DRD2-CRE line. We compared m-RNA from punched NAc of Drd2-Cre, Ankk1^lox/lox^::Drd2-Cre+ (Ankk1^Δ-DR2 Neurons^ (Ankk1^Δ-DR2 N^)) and Ankk1^lox/lox^::Drd2-Cre-(Ankk1^lox/lox^) lines, and showed a significant downregulation of *Ankk1* m-RNA in Ankk1^Δ-DR2 N^ mice as compared to Ankk1^lox/lox^ controls (**Fig.1-E and Supp.1**). However, due to the genetic construction of the DR2-CRE BAC, that bears an additional copy of Ankk1^39^, crossing leads to a downregulation of Ankk1 rather than a full knock out in DR2-expressing neurons. Interestingly, we found a downregulation of ∼24% of the *Dr2* mRNA in Ankk1^Δ-DR2 N^ mice as compared to DR2-cre controls (**Fig.1-F**), which resembles the ∼30% reduction of striatal DR2 abundance in homozygous A1 allele carriers^40^. Accordingly, with decreased DR2 activity, downregulation of Ankk1 in DR2-expressing neurons inhibited the cataleptic effects of haloperidol in Ankk1^Δ-DR2 N^ mice (**Fig.1-G**). These results suggest that the A1 variant in humans might be associated with a loss of function for the ANKK1 protein.

### Specific invalidation of Ankk1 in DR2 neurons impacts the integrative properties of DR2-SPNs

To assess the functional consequences of Ankk1 downregulation in DR2-SPNs of the NAc, we performed whole-cell patch clamp recording in brain slices of Ankk1^Δ-DR2 N^ and DR2-cre mice locally injected with a cre-dependent m-Cherry virus. DR2-SPNs neurons were identified based on tdTomato fluorescence (**Fig.2-A**). Ankk1 downregulation did not affect basic membrane properties in DR2-SPNs, as shown by resting membrane potential, resistance and rheobase (**Fig.2-A-B**, **Supp. Table1**). Furthermore, downregulation of Ankk1 in DR2-expressing neurons did not affect amplitude and frequency of spontaneous excitatory postsynaptic currents (sEPSCs, **Fig.2-C**) and spontaneous inhibitory postsynaptic currents (sIPSCs, **Fig. 2-D**). By contrast, paired stimulation (50 ms interval) of excitatory inputs onto DR2-SPNs was reduced in Ankk1^Δ-DR2 N^ mice, compared to DR2-cre mice (**Fig.2-E**). These latter data suggest that the presynaptic release probability of glutamatergic inputs onto DR2-SPNs is enhanced^41^. To further evaluate the consequences of such changes on the integrative properties of DR2-SPNs, we assessed their firing probability in response to electrical stimulation of the corpus callosum rostral to the NAc, that mostly contains glutamatergic afferents^42^. Spiking probability was significantly enhanced in DR2 SPNs of Ankk1^Δ-DR2 N^ mice (**Fig.2-F**), reflecting an increase in excitability. Overall, these data demonstrate that Ankk1 downregulation in DR2-expressing neurons enhances glutamatergic transmission onto DR2-SPNs leading to increased excitability^43^.

**Figure 1:**
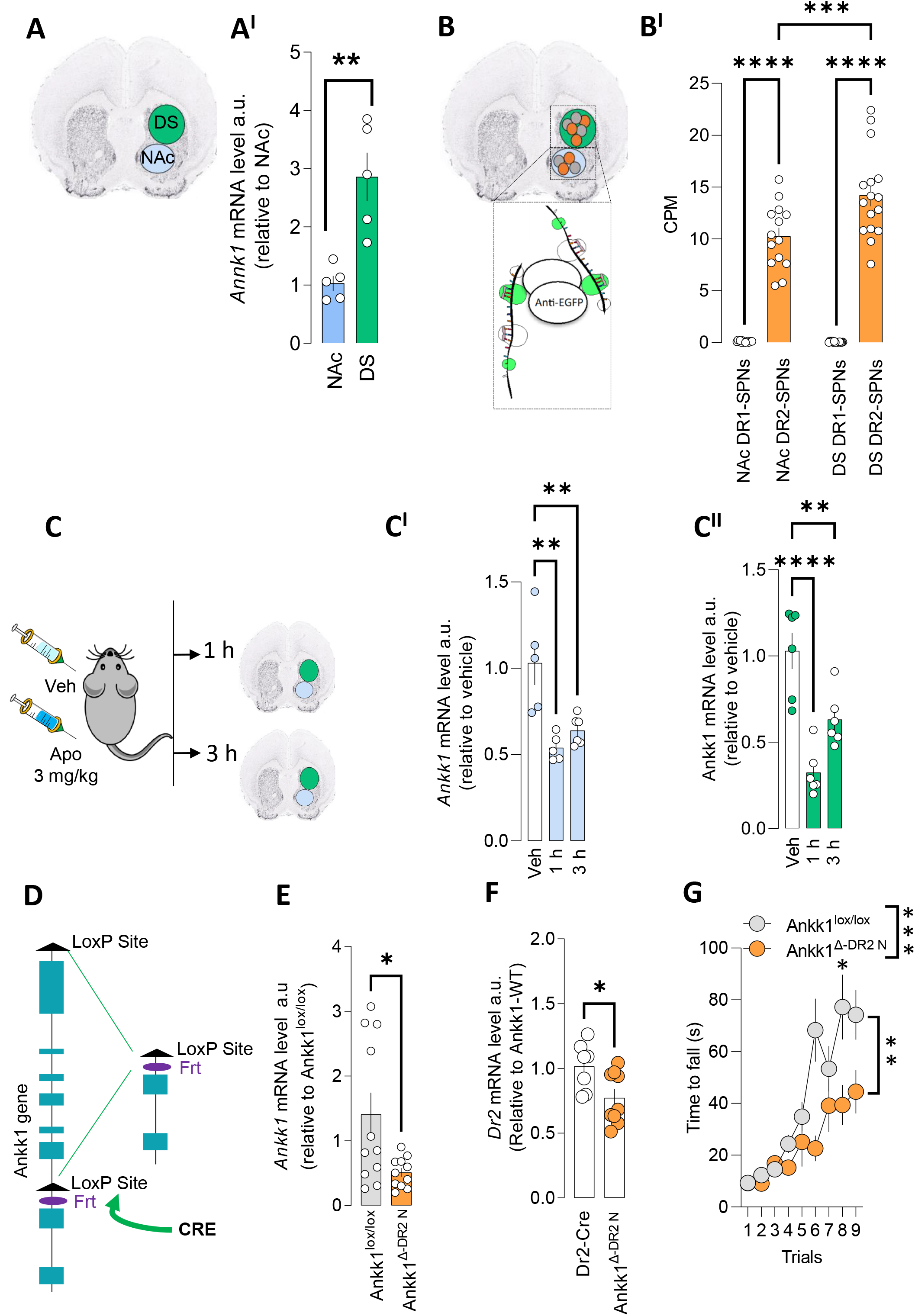
Ankk1 m-RNA is expressed in the D2-SPNs of the DS and NAc and its downregulation lead to a decrease of Dr2 m-RNA. **(A)** mRNA was purified from NAc and DS of BL-6 mice and analyzed by qRT-PCR. The expression levels were calculated by the comparative ddCt method with RPL19 as an internal control. Data points are individual results from different mice (n=5 per group). Means ± SEM are indicated. Statistical analyses are performed with two-tailed Mann-Whitney’s test, **p=0.0079. **(B)** Translating ribosome affinity purification (TRAP) technique^38^ reveals a specific enrichment of Ankk1 mRNA in DR2-SPNs as compared to DR1-SPNs in both NAc and DS. Count per millions in each immunoprecipitation are reported. Data points are individual results from different pools of mice. Means ± SEM are indicated. Two-way ANOVA Interaction F(1,56)=8,643, P=0.0048, NA/DS=F(1,56)= 8,25, P=0.0057, D1/D2 SPNs= F(1,56)=319,5-P<0.0001 Tukey’s post hoc test ****p<0.0001, *** p=0.0007. **(C)** m-RNA level of *Ankk1* in the NAc **(C**^**I**^) and the DS **(C**^**II**^) of Bl-6 mice injected with either saline or apomorphine (3 mg/Kg) and sacrificed 1h and 3h after injections. Data points are individual results from different mice (n= 5-6 per group). Means ± SEM are indicated. **(C**^**I**^) NAc, One-way ANOVA, F=11.47 p=0.0013. Dunnett’s multiple comparison p=0.0012, and p=0.0045. **C**^**II**^. DS, One-way ANOVA, F=20,87 p<0,0001. Dunnett’s multiple comparison ****p<0,0001, **p=.0046. **D**. Strategy of production of ankk1 floxed mice. **(E)** Ankk1 m-RNA level from NAc of Ankk1^lox/lox^ and Ankk1^Δ-DR2 N^ mice. Data are reported as means ± SEM of results of individual mice (n=11), p=0.0473. **(F)** Dr2 m-RNA level from NAc of Ankk1^lox/lox^ and Ankk1^Δ-DR2 N^ mice. Data are reported as means ± SEM of results of individual mice (n=7-9). Statistical analyses are performed with non-parametric Mann-Whitney test, *Dr2* p=0.0418. **(G)** The effects of *Ankk1* and *Dr2* m-RNA downregulation on DR2 function were investigated by evaluating the immobility 45-180 min after haloperidol injection (0.1 mg.kg^-1^, i.p.). Two-way ANOVA, (18 mice per group). Interaction p=0.0008, genotype p=0.0020, post-hoc Sidak test, line 6 p=0.018.

**Figure 2:**
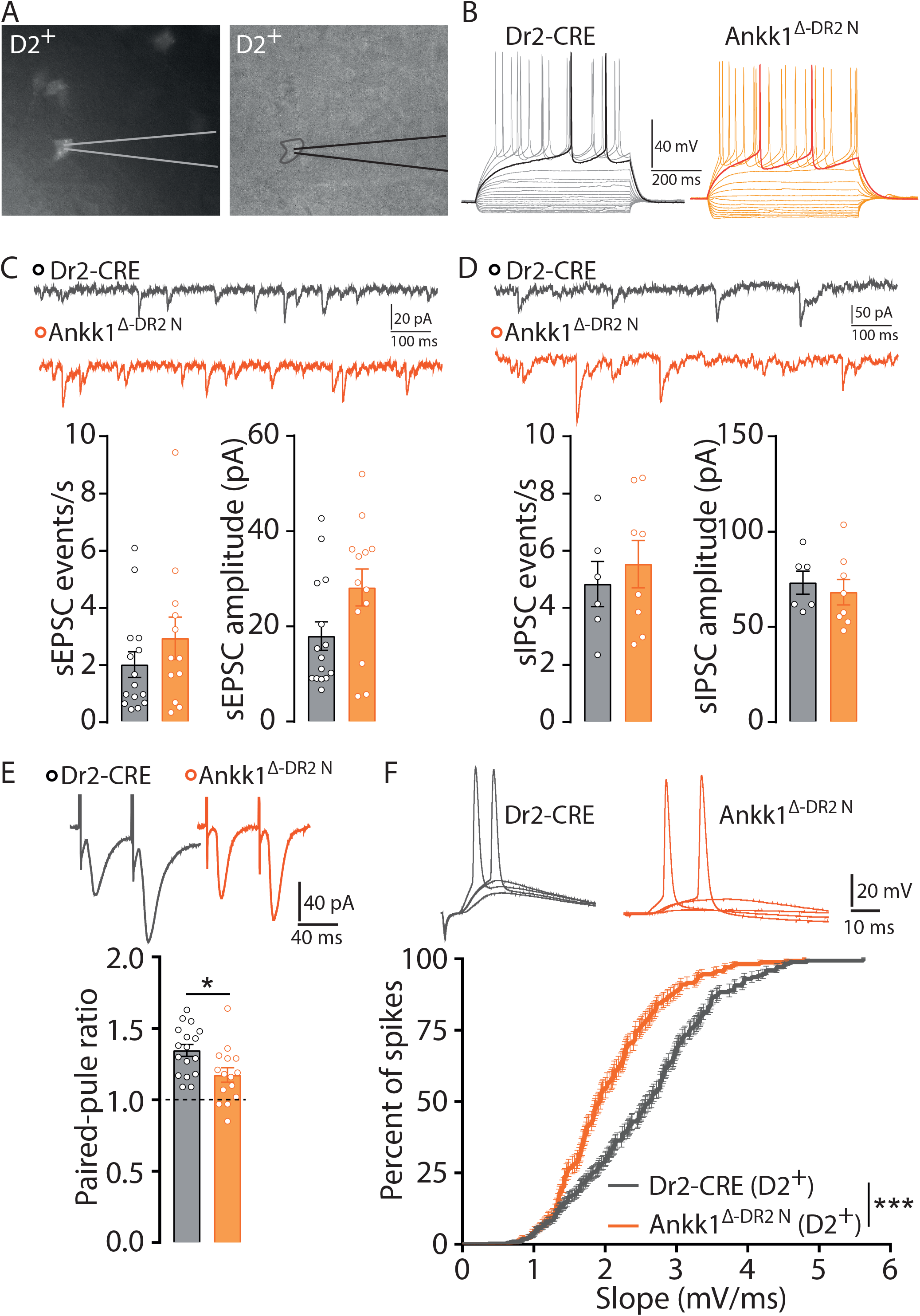
Ankk1 downregulation in DR2-expressing neurons increases their excitability. **(A)** Males and females Dr2-cre and Ankk1^Δ-DR2 N^ mice were stereotaxically injected with a viral vector carrying a flex AAV-mcherry. DR2-SPNs in the NAc were identified based on red fluorescence and patched. (**B**) Illustrative voltage responses of DR2-SPNs recorded in Dr2-cre (grey, left) and Ankk1^Δ-DR2 N^ (orange, right) mice in response to series of 600ms current pulses starting at -150 pA with 20 pA increment. **(C)** Ankk1 downregulation does not alter frequency (left) or amplitude (right) of spontaneous EPSC (sEPSC). Mann Whitney unpaired T-Test p=0.39 and p=0.09 respectively (Drd-cre: n=15 neurons in 9 mice, Ankk1^Δ-DR2 N^: n=14 neurons in 9 mice). (**D)** Ankk1 downregulation does not alter frequency (left) or amplitude (right) of spontaneous IPSC (sIPSC). Mann-Whitney unpaired T-Test p=0.59 (Dr2-cre: n=6 neurons in 3 mice, Ankk1^Δ-DR2 N^: n=8 neurons in 3 mice). (**E)** In paired pulse ratio (PPR) experiments, excitatory fibers were stimulated twice with an interval of 50ms, while EPSC were monitored in voltage clamp. In DR2-SPNs, Ankk1 downregulation resulted in a decrease of PPR. Two-tailed Mann-Whitney unpaired T-Test, p=0.0171 (Drd2-cre: n=17 neurons in 10 mice, Ankk1^Δ-DR2 N^: n=15 neurons in 9 mice). (**F)** The excitability of DR2-SPNs in the NAc was measured by quantifying the spiking probability with increasing electrical stimulation of excitatory inputs. The spiking probability is represented as a function of EPSP slope (mV/ms). Excitability of DR2-SPNs is increased in Ankk1^Δ-DR2 N^ mice. p<0.0001 (Dr2-cre: n=18 neurons in 12 mice, Ankk1^Δ-DR2 N^: n=10 neurons in 8 mice).

### Effect of Ankk1 downregulation in DR2-expressing neurons in striatal-related learning

Several studies have associated the TaqIA variant with alterations of striatal-dependent procedural learning^44^. Therefore, we next assessed the effect of Ankk1 downregulation in tasks known to strongly depend on striatal integrity. We first used striatal-dependent procedural learning based on an egocentric strategy to learn to locate, without any external cues, the baited arm in a T-maze (**Fig.3-A**, **right drawing**)^45,46,47^. Strikingly, as compared to controls, Ankk1^Δ-DR2 N^ mice displayed a strong impairment in the ability to learn the location of the reward in the maze (**Fig3-A**) supporting a key role of Ankk1 in procedural learning. We next investigated performance of controls and Ankk1^Δ-DR2 N^ mice in an operant conditioning paradigm (**Fig.3-B**, **right drawing**), another striatal-related associative task. Behavioral analyses from the operant conditioning paradigm show similar learning performances between Ankk1^lox/lox^ and Ankk1^Δ-DR2 N^ when analyzing the percentage of lever presses on the reinforced lever (**Supp.2-A**). Interestingly, although Ankk1^Δ-DR2 N^ displayed enhanced lever pressing on the reinforced lever when operant ratios were increased (**Fig.3-B**), although this result supported enhanced motivational component in Ankk1^Δ-DR2 N^ mice, performance was similar to controls on a progressive ratio task (**Supp.2-B**) where willingness to exert effort is used to measure motivation^48,49^.

Altogether, our findings show that downregulation of Ankk1 in DR2-expressing neurons leads to impairments in procedural learning, together with an alteration in reward processing; two behavioral dimensions that are main features of Taq1 carriers^44,50,51^ and strongly rely on the integrity of the striatum. Considering the emerging role of the striatum in the regulation of food intake and metabolism^52^, together with the relationship between Taq1A polymorphisms and eating behavior, we next asked whether the downregulation of Ankk1 in DR2-expressing neurons is sufficient to alter feeding patterns and whole body metabolism.

**Figure 3:**
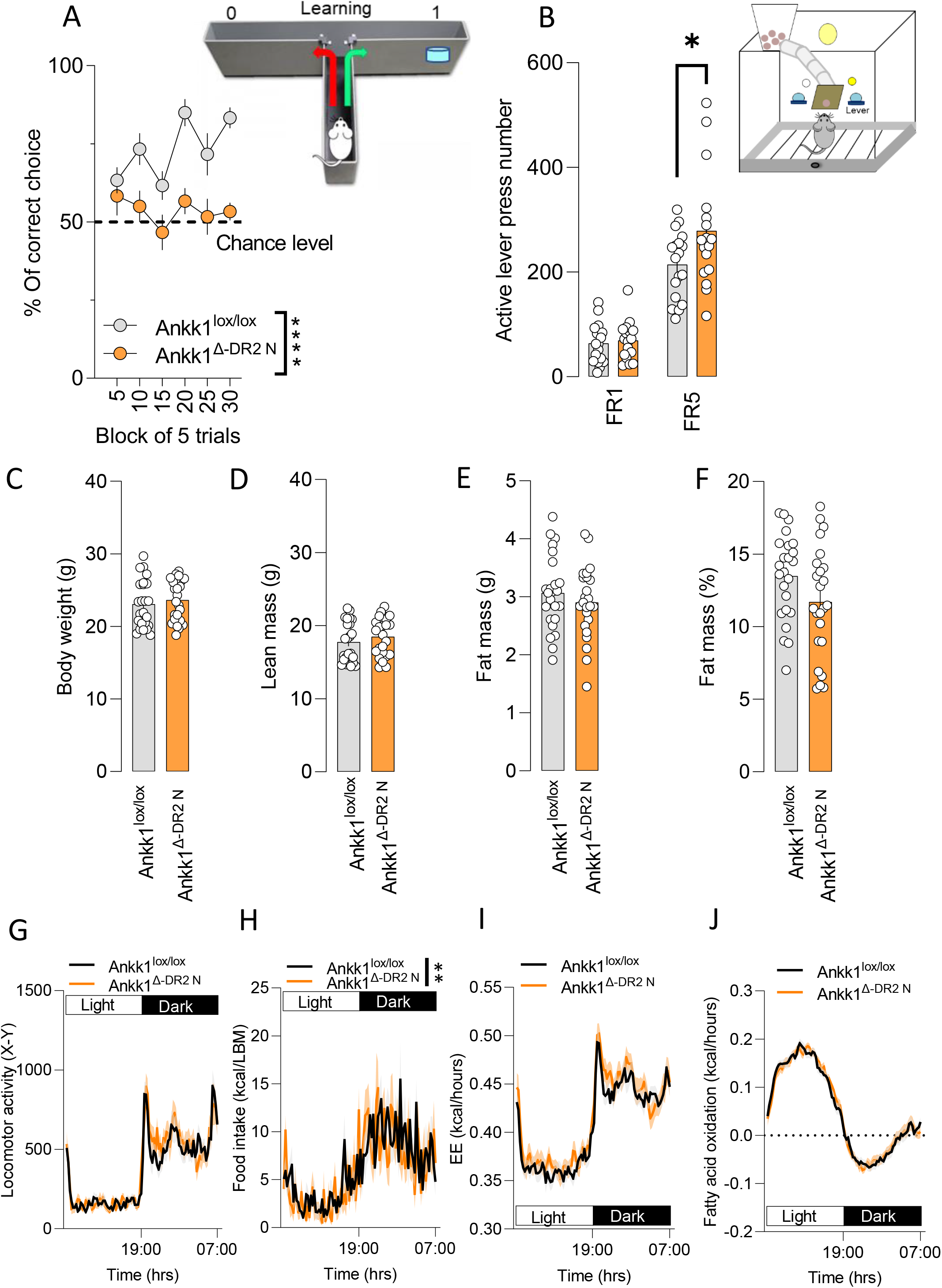
Consequences of Ankk1 downregulation in DR2-expressing neurons on striatal behavior and energy metabolism. (**A)** Effect of downregulation of Ankk1 on procedural learning. Acquisition of the food-rewarded arm choice in a T maze is impaired in Ankk1^Δ-DR2 N^ mice. Statistics: Two-way ANOVA, Interaction P=0.12, Genotype p<0.0001. N=12 (**B**) Average of active lever press per session during each period of conditioning, Two-way ANOVA, Interaction p=0.0115, Genotype p=0.0964, Time (Learning) p<0.0001. Sidak’s multiple comparison *=0.014 n=24. Ankk1 Ankk1^Δ-DR2 N^ and controls show comparable body weight (**C**), lean mass (**D**), fat mass (**E**), fat % (**F**), locomotor activity (**G**), food intake (**H**), energy expenditure **(I)**, and fatty acid oxidation **(J)**. Data are expressed as mean ± SEM. n=24 mice in each group.

### Ankk1 loss of function in DR2-expressing neurons does not alter energy homeostasis on a regular chow diet

We used indirect calorimetry to analyze metabolic efficiency and selective carbohydrates *versus* lipids substrate utilization in Ankk1^lox/lox^ and Ankk1^Δ-DR2 N^ mice fed ad libitum with a standard diet. We also measured locomotor activity and food intake. Mice showed comparable body weight (**Fig.3-C**), lean mass (**Fig.3-D**) and fat mass (**Fig.3-E**); however, there was a trend for a decrease in fat mass % in Ankk1^Δ-DR2 N^ mice (p=0.076) (**Fig.3-F**). Longitudinal measurements of metabolic efficiency and energy intake showed no differences between genotypes on locomotor activity (**Fig.3-G**), food intake (**Fig.3-H**), energy expenditure (EE) (**Fig.3-I**), or whole-body fatty acid oxidation (**Fig.3-J**).

### DR2-neurons specific Ankk1 knock down leads to change in nutrient partitioning and protection from diet-induced obesity

Indirect calorimetry investigation in Ankk1^lox/lox^ and Ankk1^Δ-DR2 N^ mice showed no substantial differences in energy homeostasis on standard chow diet. However, different studies have associated variations of *Ankk1* with metabolic changes and more specifically with obesity^23,25,53^. Hence, we next explored whether the interaction between genotype and obesogenic environment would unveil alterations in energy homeostasis regulation. To do so we subjected Ankk1^lox/lox^ and Ankk1^Δ-DR2 N^ to 3 months of a high fat high sucrose diet (HFHS) prior to metabolic characterization. Obese mice (Ob-Ankk1^Δ-DR2 N^ and Ob-Ankk1^lox/lox^) showed comparable body weight (**Fig.4-A**) and lean mass (**Fig.4-B**), however, fat mass were decreased in Ob-Ankk1^Δ-DR2 N^ mice (**Fig.4-C, D**). Ob-Ankk1^Δ-DR2 N^ and controls mice also showed similar locomotor activity (**Fig.4-E**), caloric intake (**Fig.4-F**) and a trend towards decreased EE (p=0.12) (**Fig.4-G**). Interestingly, upon consumption of the obesogenic diet, Ankk1^Δ-DR2 N^ displayed a decreased level of fatty acid oxidation (**Fig.4-H**). Such a decrease does not seem to depend on the kilocalorie intake, as shown by the lack of correlation between food intake and fatty acid oxidation (**Fig.4-I, J**). This indicates that loss of function of Ankk1 influences nutrient partitioning when mice are exposed to obesogenic diet. Altogether these data indicate that downregulation of Ankk1 in DR2-neurons impact on both adipose storage and nutrient utilization in a diet obesity paradigm (DIO).

Our findings reveal that downregulation of Ankk1 in DR2-expressing neurons is sufficient to induce both alterations in reward processing and reward-related learning, together with metabolic vulnerability to diet-induced obesity. Considering i) the alterations of striatal activity in Taq1A carriers^32,54^ and ii) the central role of the striatum in reward processing and the regulation of food intake ^55,56,57^, we next aimed to determine the contribution of striatal Ankk1 loss-of-function to such effects. We used an AAV-CRE strategy to knock-down Ankk1 in Ankk1-lox mice in the NAc (Ankk1^Δ-NAc^) and DS (Ankk1^Δ-DS^). Controls corresponded to floxed littermates injected with an AAV-GFP virus (Ankk1^GFP-NAc^ and Ankk1^GFP-DS^). Prior to behavioral analysis, accuracy for injections site and consequent change in *Ankk1* level of were assessed by immunofluorescence (**Supp.3-A, B**) and q-RT-PCR (**Supp.3-E, f**) respectively. Interestingly, while downregulation of *Ankk1* mRNA in the DS did not significantly decrease the level *Dr*2 m-RNA (**Supp.2-C**) and did not affect haloperidol-induced catalepsy (**Supp.3-D**), similar to Ankk1^Δ-DR2 N^ mice, viral-mediated knockdown of *Ankk1* in the NAc significantly decreased *Dr*2 m-RNA levels and (**Supp.3-G**) haloperidol-induced cataleptic events (**Supp.3-G**).

**Figure 4:**
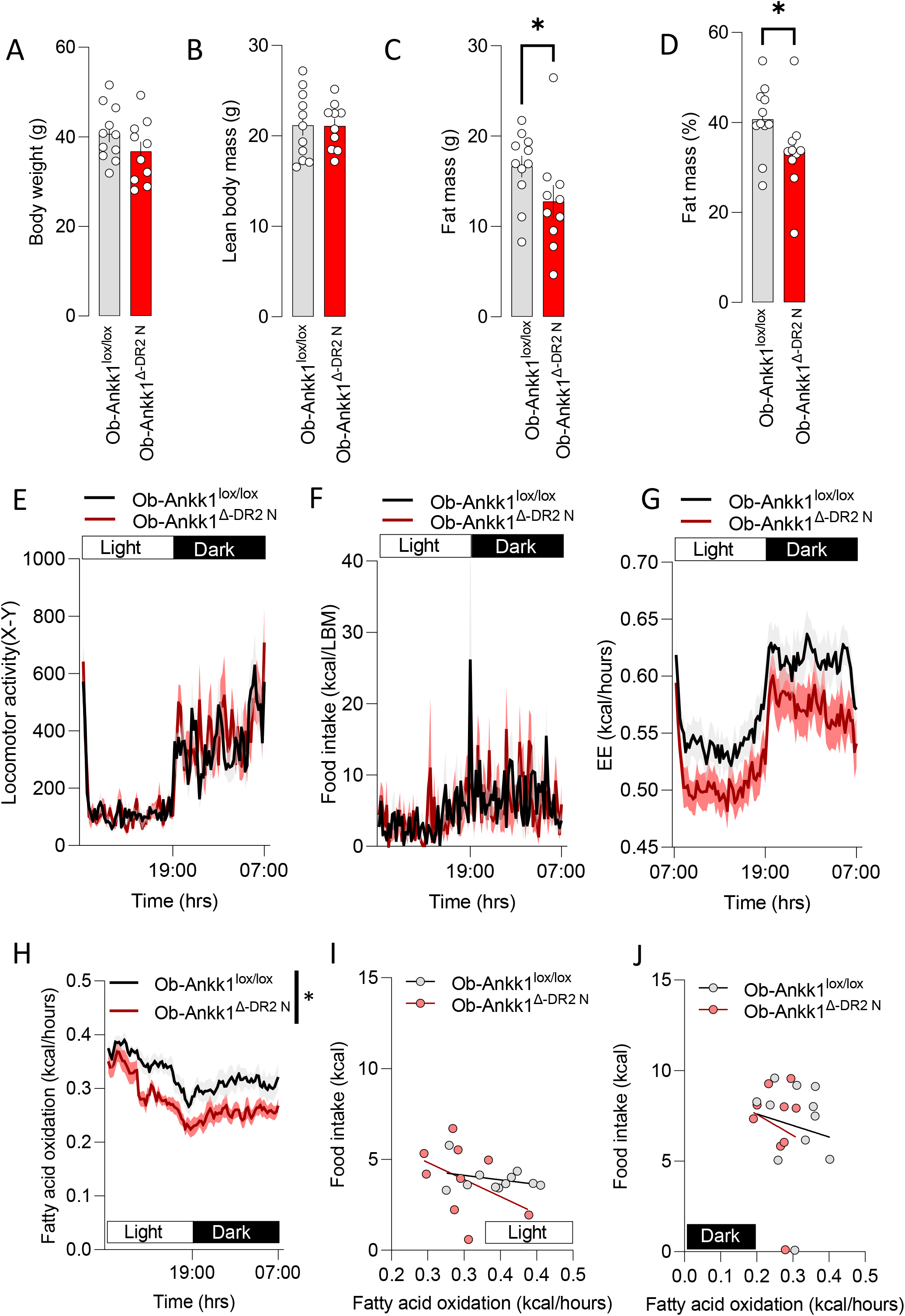
Consequences on metabolism of Ankk1 downregulation in DR2-expressing neurons in a DIO paradigm. Ankk1^Δ-DR2 N^ do not show alteration in body weight (**A**) and lean mass (**B**), however significantly decreases fat mass p=0.037 (**C**) fat mass % (**D**) p=0.0430 (Two-tailed Mann-Whitney n=11-10). Ankk1^Δ-DR2 N^ and controls show similar locomotor activity (**E**) food intake (**F**), and energy expenditure (**G**). Ankk1^Δ-DR2 N^ showed decreased fatty acid oxidation (**H**), two-way ANOVA, Interaction p=0.4487. Genotype p=0.0221, Time p<0.0001. Fatty acid oxidation does not correlate with kcal in intake in either light (**I**) or dark phase (**J**).

### Region-specific invalidation of *Ankk1* in the ventral or dorsal part of the striatum differentially affect reward-driven behavior

Interestingly, comparable to Ankk1^Δ-DR2-N^, Ankk1^Δ-NAc^ mice displayed a strong impairment in learning the egocentric strategy in the T-maze task (**Fig.5-A**), together with enhanced lever pressing already from lower ratio requirement in the operant conditioning paradigm (**Fig.5-B**). In this latter task, Ankk1^Δ-NAc^ mice exerted significantly more lever presses on the non-reinforced lever, while performance in the progressive ratio task was unchanged (**Supp.4-A, B**). Ankk1^Δ-NAc^ mice also produced significantly more lever presses on the active lever during the inactive phase, a proxy for impulsive behavior^58^. Importantly, this effect in the progressive ratio task was observed in both *ad libitum* and food restricted conditions, indicating that this phenotype is not strictly dependent ofhunger state (**Supp.4-C, D**). Together, these findings suggest that, rather than enhanced motivation, augmented performance in the operant conditioning paradigm in Ankk1^Δ-NAc^ mice might be related to increased impulsivity^58^, a main feature of Taq1A carriers^34,35,59,60^.

Altogether, our findings demonstrate that Ankk1 loss-of-function in the NAc is sufficient to recapitulate the reward-related phenotypes of Ankk1^Δ-DR2 N^ mice. By contrast, Ankk1^Δ-DS^ mice had similar performance as controls in both the learning phase of the T-maze paradigm, as well as in operant conditioning for both ratio requirements. However, Ankk1^Δ-DS^ mice displayed a selective deficit in the reversal phase of the T-maze (**Supp.5 A-B**). Of note, this latter finding resembles impaired cognitive flexibility described in Taq1A carriers^26,61^.

**Figure 5:**
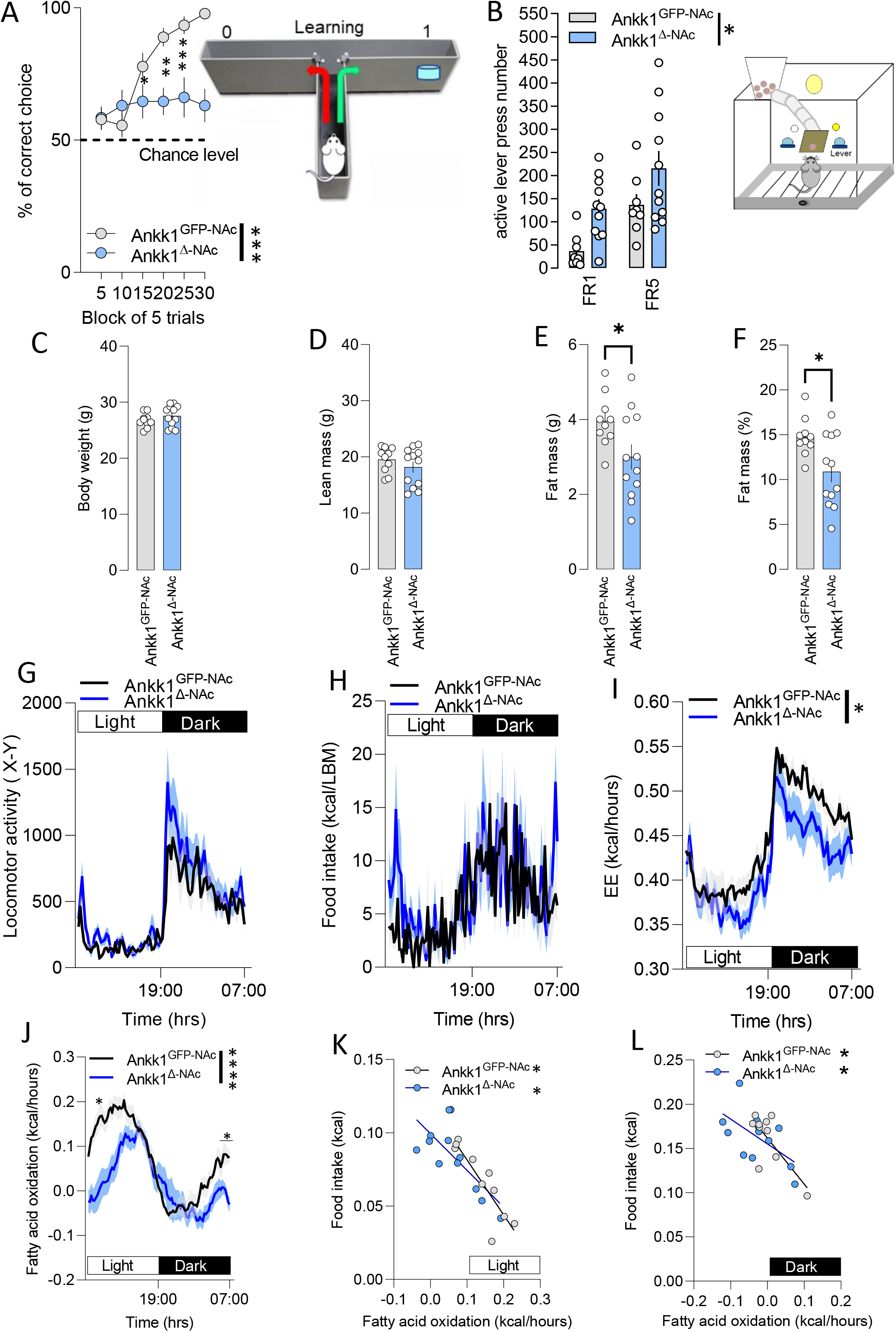
Consequences of Ankk1 downregulation in NAc on striatal-related behavior and energy metabolism. **(A)** Effect of Ankk1 loss-of-function in the NAc on procedural learning. Acquisition of the food-rewarded arm choice in a T-maze is impaired in Ankk1^Δ-NAc^ mice. Statistics: two-way ANOVA, Interaction p=0.0004, Time p<0.0001, Genotype p<0.0001. Sidak’s multiple comparison *=0,0176 **=0,0055 ***=0,0002 N=9-13. (**B**) Average of active lever press per session during each period of conditioning, is increased in Ankk1^Δ-NAc^ as compared to Ankk1^GFP-NAc^.Two-way ANOVA, Interaction p=0.776, Learning p=0.0006, Genotype p=0.0162. N=8-11 **C**. Ankk1 loss-of-function in the NAc does not alter body weight (**C**) and lean mass (**D**) but induces a decrease in fat mass (**E**) and fat mass % (**F**), Two tailed T-test, t=2,284, df=20 p=0.0335 fat mass t=2,696, df=20 p=0.0139, respectively. Data are expressed as mean ± SEM. n=10-13. Ankk1^Δ-NAc^ and Ankk1^GFP-NAc^ showed comparable locomotor activity (**G**) and food intake (**H**), however KD of Ankk1 led to a decreased energy expenditure (**I**), statistics: Two-way ANOVA, Interaction p=0,034 Time, p<0.0001, genotype= F _(1, 20)_ = 2,820 p= 0.108, and fatty acid oxidation (**J**) Two-way ANOVA, Interaction p<0.0001, p<0.0001, p=0.0075 Sidak’s post-hoc test *<0.05. Fatty acid oxidation significantly correlates with food intake for light (**K**) and dark phase (**L**). Statistics light phase CTR p=0.0009, KD, p=0.0083, dark phase Ankk1 CTR p=0.0152, KD, p=0.0441.

### Striatal deletion of Ankk1 alters energy homeostasis

We next assessed metabolic parameters in DS and NAc Ankk1 KD mice. Ankk1 loss-of-function in the NAc did not result in any significant changes of body weight (**Fig5-C**) and lean mass (**Fig5-D**). However, KD mice showed decrease fat mass (**Fig5-E**) and fat mass % (**Fig5-F**). As for the Ankk1^Δ-DR2 N^ groups, Ankk1^Δ-NAc^ mice showed unaltered locomotor activity (**Fig.5-G**) and caloric intake (**Fig.5-H**), although a trend was detected (Genotype p=0.056, Interaction p=0.06) (**Fig.5-D**). Moreover, Ankk1^Δ-NAc^ displayed a decrease EE (**Fig.5-I**) and fatty acid oxidation (**Fig.5-J**). For both groups, the variations in fatty acid oxidation correlated with caloric intake (**Fig.5-K, L**), but was independent from body weight and lean mass which are comparable between the two genotypes (**Fig.5-C, D**), indicating that loss of function of Ankk1 influences peripheral nutrient utilization. Interestingly, fatty acid oxidation mostly decreased during the light phase (**Fig.5-L**), which might indicate a dissociation between circadian-entrained-rhythm and whole body fatty acid oxidation^62,63^. Importantly, when tested on a binge eating paradigm with HFHS, a test aiming at quantifying uncontrolled voracious eating, Ankk1^Δ-NAc^ mice showed enhanced food consumption during the binge time (**Supp.4-E**). Therefore, in order to explore whether high fat diet exposure could affect feeding patterns in the Ankk1^Δ-NAc^ group, we next tested control and KD mice in a DIO paradigm. Mice were fed a HFHS diet for 3 months prior to the replication of indirect calorimetry measurements. As compared to controls (Ob-Ankk1^GFP-NAc^), obese (Ob) Ob-Ankk1^Δ-NAc^ mice also showed a decrease in body weight (**Fig.6-A**), comparable lean mass (**Fig.6-B**), a decrease in fat mass (**Fig.6-C**) and a tendency of decreasing fat mass % (**Fig.6-D**). Ob-Ankk1^Δ-NAc^ mice showed comparable caloric intake (**Fig.6-E**), higher locomotor activity (**Fig. 6-F**), and decreased energy expenditure (**Fig.6-G**). As for Ob-Ankk1^Δ-DR2 N^, Ob-Ankk1^Δ-NAc^ mice diplayedlower fatty acid oxidation (**Fig.6-H**), which was independent from food intake (**Fig.6-I, J**). Overall, these data indicate that Ankk1 loss-of-function in the NAc exert some protective effect from HFHS diet-induced disturbance in nutrient intake and partitioning. Confirming a main role of Ankk1 in the NAc, Ankk1 loss-of-function in the DS did not result in any relevant alteration in any of the measured energy homeostasis parameters. In fact, body weight, lean mass and fat storage were comparable between KD and CTR groups (**Supp.5-D, G)**. Locomotor activity (**Supp.5-H**), feeding (**Supp.5-I**) and energy expenditure (**Supp.5-J**) were also comparable between groups. However, we could observe a small difference in the light-dark phase distribution of whole fatty acid in oxidation Ankk1^Δ-DS^ mice (**Supp.5-H**). No difference between groups were observed in those mice in a binge eating paradigm (**Supp.5-C**).

**Figure 6:**
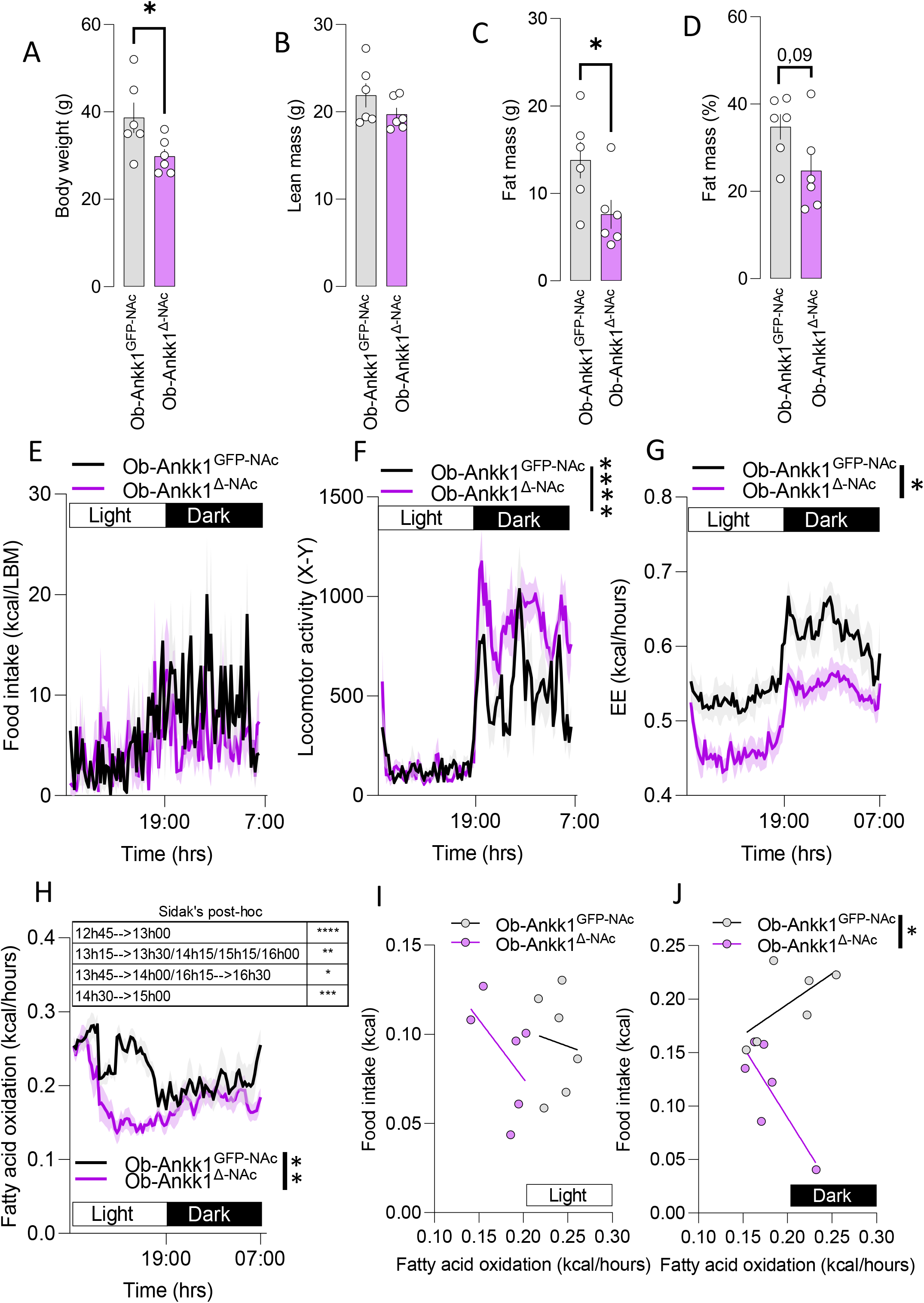
Consequences of Ankk1 loss-of-function in the NAc on metabolism in a DIO paradigm. Ankk1^Δ-NAc^ display decreased body weight (**A**)Two-tailed Mann-Whitney’s test, p = 0.0455, but not lean mass (**B**), as well as decreased fat mass (**C**) Two-tailed Mann-Whitney’s test, p=0.0411, but not fat mass percentage (**D**) Two-tailed Mann-Whitney’s test, p=0.0931. **E** caloric intake is comparable between Ankk1^GFP-NAc^ and Ankk1 ^Δ-NAc^ but the Ankk1 ^Δ-NAc^ group showed increased locomotor activity (**F**) and decreased energy expenditure (**G**). Statistics F: Two-way ANOVA, Interaction p<0,0001 Time, p<0.0001, genotype p=0.0138. Statistics G: Two-way ANOVA, Interaction p<0,0001 Time, p<0.0001, genotype p=0.0222. **H** Downregulation of Ankk1 in the NAc decreases fatty acid oxidation, statistics: Two-way ANOVA, Interaction p<0.0001, Time p<0.0001, genotype p=0.0074. Sidak’s post hoc test, *=p<0.05, **=p<0.001 ***=p<0.0001. **I-J** Fatty acid oxidation does not significantly correlate with food intake in both light and dark phase, however, in the dark phase slopes of the regression lines are significantly different between the two groups p=0,0129 (**J**). Data are expressed as mean ± SEM. n=6.

**Figure 7:**
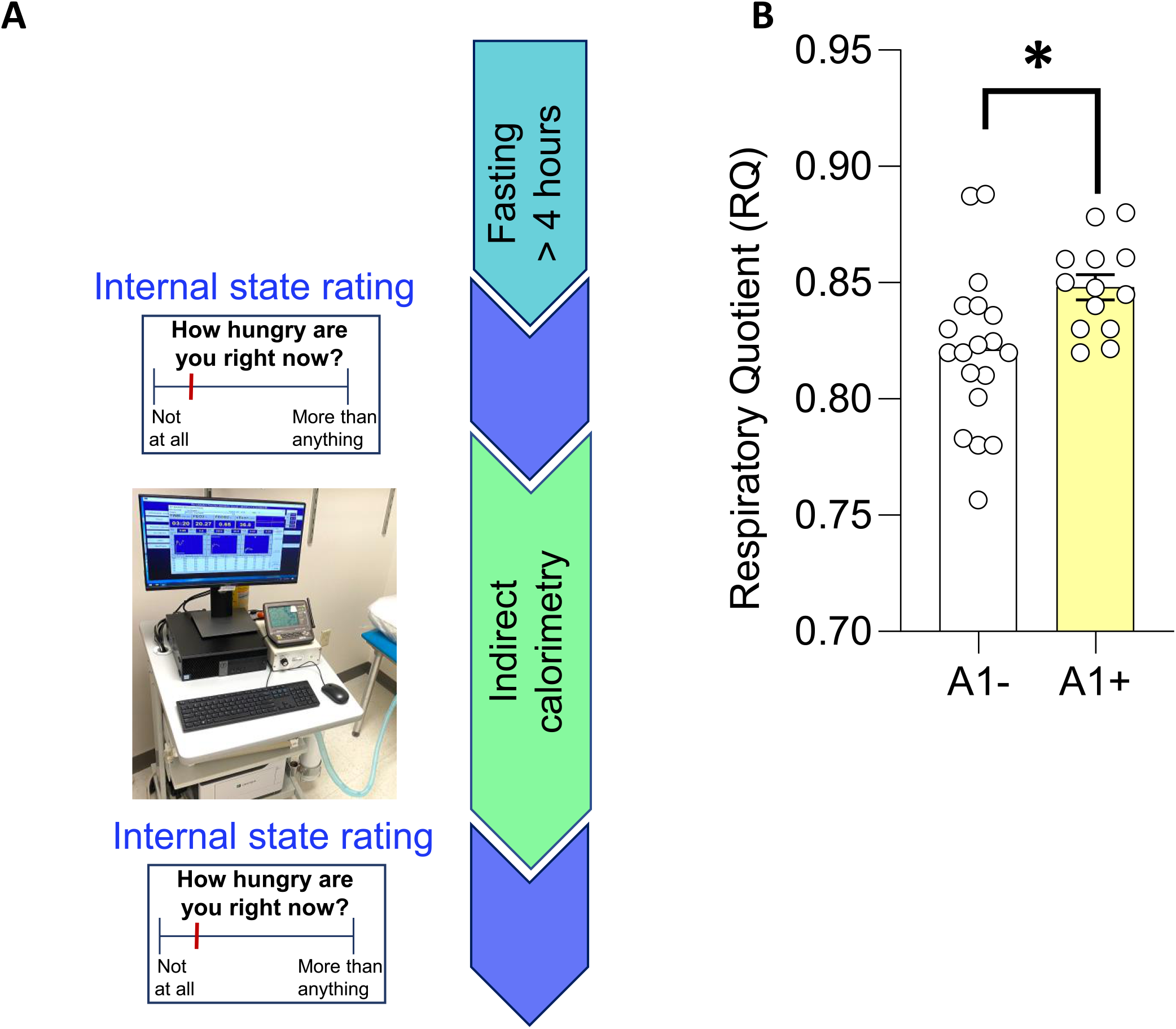
Resting respiratory quotient (RQ) between A1 allele carriers (A1+) and non-A1 carriers (A1-) of Taq1A polymorphism (rs1800497). 32 participants with healthy weight (BMI<26) underwent metabolic measurements using indirect calorimetry and genotyping from saliva. Data points are individual results from different participants (n=19 in A1- and n=13 in A1+). Means + SEM are indicated. Statistical analysis is performed with an independent t-test and controlled for BMI, sex, age, and study. *p=0.012.

### Differential respiratory quotient as a function of Taq1A A1 allele status in human participants

Given, the observed metabolic phenotype in mice, as a final step, we used indirect calorimetry during resting state to calculate the respiratory quotient (RQ) (**Fig.7**) in 32 healthy human participants (19 A1- and 13 A1+). Age (A1−: M = 27.8, SD = 1; A1+: M = 26.9, SD = 5.8; *t* = 0.29, *p* = 0.77), BMI (A1−: M = 22.3, SD = 2.4; A1+: M = 22.6, SD = 1.9; *t* = 0.35, *p* = 0.73), fat mass (A1−: M = 17.6, SD = 6; A1+: M = 14.9, SD = 4.3; *t* = 1.05, *p* = 0.30), and other anthropometric measures were similar for both genotypes (**Supp. Table 2**). The groups also did not differ in hours of sleep and hours since last meal prior to the metabolic measure (**Supp. Table** 2). However, there was a significant difference in sex distribution between A1+ and A1 groups (*p* = 0.0014) (Supp. Table 2) so this factor is included in statistical models. Consistent with the observations in mice, A1+ individuals showed significantly higher resting respiratory quotient (RQ) compared to A1-individuals (R^2^ = .41, F(6) = 7.31, p = 0.012). RQ indicates the volume of carbon dioxide released over the volume of oxygen absorbed during respiration, hence, this result suggests a shift towards carbohydrate use as a primary energy source in A1+ and a shift towards fat in A1-at rest. This indicates that polymorphism affecting Ankk1 gene in human does indeed alter peripheral nutrient utilization. This data supports our reverse translational approach and show that that the metabolic phenotype observed in mice could translateto humans.

## DISCUSSION

The elevated occurrence of metabolic dysregulations in psychiatric disorders suggests the existence of shared pathogenic pathways with neuropsychiatric symptoms^4,64^. Genome-wide association studies (GWAS) have revealed substantial genetic components in the risk for psychiatric diseases, but also for metabolic disorders^65,66^. Among all the known polymorphisms, TaqIA single nucleotide variant, located in the gene that codes for the tyrosine kinase ANKK1, bears a particular interest since it increases the risks for both metabolic syndrome as well as psychiatric disorders. From a neurobiological point of view, the TaqIA polymorphism has been associated with alterations of DA transmission within the so called reward system, in particular a decrease in DR2 binding in the striatum^54,67^, a main feature of several mental disorders such as compulsive eating^68^ or addiction^8^, whose prevalence is increased in TaqIA carriers. Strikingly, we found that *Ankk1* is enriched in DR2-SPNs and that its downregulation selectively in DR2-expressing neurons is sufficient to decrease *Dr2* mRNA expression. These data are in line with previous work in humans indicating that such downregulation in psychopathologies would take place at the transcriptional level^15^. DR2 expression is known to play a key role in the functionality of DR2-SPNs^69,70,71^. Even though it is unlikely that these effects are only related to decreased DR2 expression, we found that downregulation of Ankk1 in DR2-expressing neurons leads to increased excitability of DR2-SPNs.

Increased DR2-SPNs activity is consistent with impaired reward-related behaviors^71,72,73^, in particular regarding the NAc, considering its central role in reward processing^74,75^. In fact, viral-mediated deletion of Ankk1 selectively in the NAc at adulthood recapitulate and amplifies some of the behavioral phenotypes obtained in Ankk1^Δ-DR2 N^, such as alteration in procedural learning and operant behavior. DR2 being expressed in different central and peripheral areas, a specific effect of Ankk1 KD in DR2-expressing cells of other brain areas, such as ventro tegmental area, cannot be excluded. Also, differences in phenotypes strength might originate from the developmental effects in Ankk1^Δ-DR2 N^ mice, which could lead to compensatory mechanisms that could temper some of the phenotype observed in Ankk1^Δ-NAc^. The consistent deficits we observed in the T-maze task for both Ankk1^Δ-DR2 N^ and Ankk1^Δ-NAc^ are unlikely to be simply related to learning inabilities. In fact, manipulations of the NAc can spare the acquisition of action-outcome associations, while impairing flexible adaptation of previously learned rules^76,77,78^, in particular when interfering with DR2-SPNs^79^. In accordance, both Ankk1^Δ-DR2 N^ and Ankk1^Δ-NAc^ were capable of learning the association between lever pressing and reward obtainment. However, Ankk1^Δ-NAc^ performed significantly more lever presses on the non-reinforced lever, suggesting increased impulsivity^58^. Such a behavioral alteration might interfere with the acquisition of the egocentric strategy in the T-maze and could explain why both Ankk1^Δ-DR2 N^ and Ankk1^Δ-NAc^ lines display higher performance for low ratio requirement in the operant conditioning paradigm, but not in the motivation-based progressive ratio task. Interestingly, impulsivity has been related to decreased DR2 availability in the NAc^80,81,82^, further implicating NAc DR2-SPNs in the behavioral effects of Ankk1 loss-of-function. Importantly, all the aforementioned behavioral effects are in line with the A1 allele being associated with poorer negative outcome learning^50,51^ and procedural learning^44^, increased impulsivity^34,35,59,60^, and more generally altered prediction error^50^ and weaker reward sensitivity (for a review^37^). Of note, the selective impairment in the reversal phase of the T-maze in Ankk1^Δ-DS^ resembles decreased cognitive flexibility^26,61^ and reduced activity in the prefrontal cortex and striatum during reversal learning^26,50^ in Taq1A carriers. Overall, our findings support that behavioral deficits related to Ankk1 loss-of-function mostly originate from dysregulation of DR2-SPNs of the NAc. This is in line with the stronger reduction of DR2 binding in the ventral striatum compared to the dorsal striatum in TaqI1 carriers^54^.

In addition to recapitulating reward-related behavioral alterations characteristic of Taq1A carriers, Ankk1 loss-of-function in the NAc was also sufficient to impact whole energy homeostasis. In fact, Ankk1^Δ-NAc^ displayed decreased fatty acid oxidation. Comparable to the behavioral effects, alterations in lipid metabolism were strengthened in Ankk1^Δ-NAc^ mice as compared to Ankk1^Δ-DR2 N^. Interestingly, the main metabolic features seemed spared under regular diet while specifically triggered on obesogenic diet in which Ankk1 ^Δ-DR2-N^ mice displayed metabolic alterations similar to those observed in both lean and Ob-Ankk1^Δ-NAc^ mice.

Specifically, in both models, Ankk1 loss-of-function led to decreased fatty acid oxidation and was protective against fat mass increase during DIO. This suggests that an interaction between genotype and obesogenic environment is necessary to unmask metabolic alterations in energy homeostasis in Ankk1^Δ-DR2-N^ mice and is in line with recent findings highlighting a particular sensitivity of DR2-SPNs to high fat diet and circulating lipids^11,10,42,83^. This observation is particularly relevant from a translational point of view since, in humans, genetic polymorphisms, including Taq1A, represent a risk factor for pathological conditions, in interaction with the environment. Our findings demonstrate an intriguing role of the NAc in the regulation of metabolism that goes beyond its established implication in the regulation of feeding behavior, through its connections with the lateral hypothalamus and the l ventral tegmental area^55,56,57,84^. Importantly, while vulnerability to the development of metabolic syndrome in Taq1A carriers has been largely related to alterations in reward (palatable food)-oriented behavior, our findings suggest that a direct brain-mediated dysregulation of metabolism might also be at play. Consistent with this possibility, we also report preliminary evidence that human carriers of the A1 allele show decreased fatty acid oxidation at rest compared to individuals without the allele.

However, rather than favoring the development of obesity-like phenotypes, Ankk1 loss-of-function seems to protect from increases in body weight and fat mass. Yet, as previously mentioned the A1 polymorphism has also been associated with accelerated weight loss^23^ and some features of anorexia^22^, which share common symptomatic dimensions with compulsive eating such as deficit in cognitive flexibility, impaired reward processing and impulsivity^85, 86^. It is thus tempting to propose that vulnerability to the development of obesity in Taq1A carriers might be mostly related to reward-related alterations leading to compulsive eating that would overcome metabolic regulation. Alternatively, it is also possible that the metabolic features we observed under Ankk1 loss-of-function reflect a higher vulnerability of peripheral metabolism that could lead, on the long term, to opposite phenotypes depending on dysregulation of food intake. Even though highly speculative, such a hypothesis might explain why the very same polymorphism might render vulnerable to two opposite eating disorders, i.e. obesity or anorexia. Our study also offers a potential rational regarding the observation found in human of an interaction between TaqIA/Ankk1 polymorphism and the fat mass and obesity-associated gene (FTO)-the strongest common genetic determinants of adiposity-with increasing risk of metabolic^25^. Future work using both rodent and humans are needed to explore if and how the structure of the Ser/Thr kinase ANKK1 could be exploited in its A1/A2 variant to promote drug engineering for the treatment of altered feeding behavior and body weight gain.

In conclusion, we found that Ankk1 loss-of-function in mice, by dysregulating DR2-SPN, mostly in the NAc, mimics the main reward-related as well as metabolic symptomatic dimensions characteristic of Taq1A carriers. Importantly, even though the underlying mechanisms remain to be established, Ankk1 loss-of-function is sufficient to recapitulate the decrease in DR2 expression characteristic of Taq1A polymorphism and that is believed to be at the core of the vulnerability to develop psychiatric and eating disorders. It has been shown that the amount of physical activity correlates with DR2 expression level in the striatum, and that increased physical activity positively correlates with ameliorated cognitive functions. Hence, it is tempting to hypothesize that the behavioral alterations caused by Ankk1 loss of function would benefit from a combination of diet and physical activity that, by decreasing the level of circulating and stored lipids, would amplify the weight of metabolic alterations we observed in these mice models.

## Supporting information

Supplemental Statistic information

## AUTHORS CONTRIBUTION

**EM**, conceived the project, designed the experiments, analyzed and interpreted the data, wrote the manuscript. **RW**, performed the electrophysiology experiments. **JC** performed the NAc surgeries, **EF** performed surgeries in DS, **AA, RH, AP, JB** performed experiments. **JB, ACS** Edited the manuscript **XF, ZH, SM** performed human experiments. **GG** discussed the data and provided input to experiments and MS, **CM** supervised fiber photometry experiments, **PT** supervised the electrophysiology experiments, provided input and improvements to behavioral experiments and MS **CBB** supervised and performed the electrophysiology experiments, **D.M.S** conceived and supervised the clinical part of the project provided input and corrections to the manuscript. **SL** conceived and supervised the overall project, analyzed and interpreted the data, secured funding provided input and corrections to the manuscript.

### ACKNOWLEDGEMENTS

This work was funded by FRM Project #EQU202003010155. We acknowledge funding supports from the Modern Diet and Physiology Research Center (MDPRC), the Centre National de la Recherche Scientifique (CNRS) through the International Research Project (IRP) « BrainHealth », L’agence Nationale de la Recherche (ANR) ANR-19-CE37-0020-02, The Université Paris Cité, INRAE, Université de Bordeaux, the « Fédération pour la recherche sur le cerveau (FRC) » and the « Fondation des Treilles » « Fondation des Treilles créée par Anne Gruner Schlumberger, a notamment pour vocation d’ouvrir et de nourrir le dialogue entre les sciences et les arts afin de faire progresser la création et la recherche contemporaines. Elle accueille également des chercheurs et des écrivains dans le domaine des Treilles (Var) www.les-treilles.com » EM was supported by a post doctoral fellowship from the FRM. We thank Professor Ralph DiLeone for inputs on the study. We thank Olja Kacanski for administrative support, Isabelle Le Parco, Aurélie Djemat, Daniel Quintas, Magguy Boa and Ludovic Maingault and Angélique Dauvin for animals’ care and Florianne Michel for genotyping. We acknowledge the technical platform Functional and Physiological Exploration platform (FPE) of the Université Paris Cité, CNRS, Unité de Biologie Fonctionnelle et Adaptative, F-75013 Paris, France, the viral production facility of the UMR INSERM 1089 and the animal core facility “Buffon” of the Université Paris Cité/Institut Jacques Monod. We thank the animal facility of IBPS of Sorbonne Université, Paris.

## MATERIAL AND METHODS

### Animals

C57BL/6-Ankk1^*tm1aNarl*^ (Strain name: RMRC13296: Ankk1 Ankk1^FrtNeoLacZ/+^) were obtained from the National Laboratory Animal Center (NLAC). Ankk1^FrtNeoLacZ/+^ result from the insertion of a FRT*-En2* Intron-*En2* Exon-IRES-LacZ-***LoxP***-PGK-Neo FRT:***LoxP*** between exon 2 and 3 in addition to an insertion of a LoxP site after exon 8. Ankk1^FrtNeoLacZ/+^ were bred with B6N.129S4-Gt(ROSA)26Sor*tm1(FLP1)Dym/J (Rosa26::FLPe Knock in)* expressing the flippase under the ubiquitous Rosa26 promoter. The resulting mice Ankk1^*lox/lox*^ mice contains a FRT-LoxP site between Exon 2 and 3 in addition to an insertion of a LoxP site after exon 8. Cre-mediated recombination result in the excision of Exon CRE-mediated recombination 3-8, ENSMUSE00000218187-ENSMUSE00000408787 (Supplementary Figure 1). Ankk1^*lox/lox*^ mice are genotyped using primer Ankk1LF (64) 5’-AAT GCC GCT GAG CAG TCA GGC TGG-3’ and Ankk1LR (65) 5’-GCC TAG TGA AGG CTC CTT GTC CTG-3’ which produces a 300bp in WT animals and a 440bp in floxed Ankk1 animals, (PCR protocol: step 1: 94°C,3min. Step 2: (35 cycles) 94°C-20 sec::60°C-15sec::72°C-30sec. Step 3: 72°C 2 min).

Ankk1^Δ-DR2 N^ transgenic animals were produced by breading Annk1^lox/lox^ and Dr2-Cre animals and maintained on a C57Bl/6J background. Drd2-Cre animals were from Genesat Drd2-Cre (Tg(Drd2-cre)ER44Gsat/Mmucd). Wilt-type C57Bl/6 mice were purchased from Janvier (France) and used at 10-12 weeks. Male and females were used for experiments. Mice were maintained on a 12-h light/dark cycle (light on at 7:00 am) and had, before the beginning of the experiment, free access to water and food. Animal were fed with either chow diet or high-fat high-sugar diet (HFHS, cat n. D12451, Research Diets, New Brunswick, USA) for twelve to sixteen weeks. Animal protocols were performed in accordance with the regulations and approved by the relevant committee: Paris, guidelines of the French Agriculture and Forestry Ministry for handling animals (decree 87-848) under the approval of the “*Direction Départementale de la Protection des Populations de Paris*” (authorization number C-75-828, license B75-05-22), Animal Care Committee of the University of Paris (APAFIS # 2015062611174320), institut de Biologie Paris Seine of Sorbonne University (C75-05-24).

### Total RNA purification, cDNA preparation and real-time PCR

Mice were sacrificed by decapitation. The brain was quickly dissected out, placed in cold buffer and then in an ice-cold brain form to cut thick slices from the nucleus accumbens (NAc) and the dorsal striatum (DS) punched out using ice-cold stainless steel cannulas. Tissue samples (1 mouse per sample, dissected as above) were homogenized in TRIzol with loose and tight glass-glass 2-mL Dounce homogenizers. Total RNA was extracted with TRIzol Reagent (Life Technologies) according to manufacturer’s instructions. The RNA was quantified with a Nanodrop 1000 spectrophotometer. RT-qPCR, was performed using SYBR Green PCR kit in 96-well plates according to the manufacturer’s instructions. 500 ng of RNA were used for reverse-transcription, performed with the MMLV reverse transcriptase (Invitrogen). qPCR was performed in a LightCycler 1.5 detection system (Roche, Meylan France) using the LightCycler FastStart DNA Master plus SYBR Green I kit (Roche). Results were quantified and normalized to a house-keeping gene (RPL19) with delta-delta-CT (ddCT) method.

### Pharmacological treatments

For acute treatments, apomorphine (Tocris) was dissolved in phosphate-buffered saline (PBS) and injected i.p. (3 mg.kg^-1^). PBS was used as vehicle treatment in control mice. Haloperidol (Tocris) was dissolved in saline and injected i.p. (0.5 mg.kg^-1^).

### Histology

Ankk1^GFP-NAc^, Ankk1^Δ-NAc^, and Ankk1^GFP-DS^ Ankk1^Δ-DS^ mice were transcardially perfused with 40 g.L^-1^ paraformaldehyde (PFA) and post-fixed overnight. Coronal sections (30 µm) were prepared on a cryostate. Sections were blocked with 25 g.L^-1^ bovine serum albumin (BSA) in PBS containing 0.2% Triton. Afterwards, slices were incubated overnight with at 4 C° with a chicken anti-GFP primary antibody (1:2000, ABCAM) mixed in the same blocking solution. Subsequently, slices were rinsed in PBS 3 times for 10 min each at room temperature, after which they were incubated with a corresponding secondary antibody Alexa Fluor 488 (1:500, Life Technologies) for 2 h. Slices were again rinsed 3 times for 10 min in PBS and placed in a PBS-DAPI solution for 10 min. Afterwards, slices were rinsed a final time for 10 min in PBS and then mounted and coverslipped using DAPCO adhesive. Image acquisition was performed with a Confocal system (Zeiss) using x 10 and x 40 objectives.

### Patch-clamp recordings of DR2-SPNs neurons in the NAc

Slices containing the NAc were prepared from males and females Dr2-cre and Ankk1^Δ-DR2N^ mice. After Lurocaïne (30 mg/kg) and Exagon (300 mg/kg) lethal injection, an intra-cardiac perfusion was made with N-Methyl-D-glucamine (NMDG) solution containing (in mM): 1.25 NaH2PO4-H2O, 2.5 KCl, 7 MgCl2-H2O, 20 Hepes, 0.5 CaCl2-2H2O, 28 NaHCO3, 8 D-Glucose, 5 L(+)-Ascobate, 3 Na-Pyruvate 93 NMDG, bubbled with carbogen gas and pH adjusted to 7.35 with HCl. The brain was quickly removed from the skull and sagittal slices (350 µm) were obtained with a vibratome (VT1000S, Leica) in cold NMDG solution. After slicing, slices were immediately transferred in NMDG solution at 34°C for 12 min. Then, slices were transferred individually at room temperature in artificial cerebrospinal fluid (ACSF) containing (in mM): 125 NaCl, 25 NaHCO3, 2.5 KCl, 1.25 NaH2PO4, 2 CaCl2, 1 MgCl2, 25 glucose and 1.25 pyruvate, bubbled with carbogen gas. Slices were used after at least 1 hour of rest. Recordings were performed as previously described^42^. Briefly, slices were recorded at 30°C in a recording chamber continuously superfused at 1-1.5 ml/min with oxygenated ACSF. Neurons were visualized under an upright fluorescent microscope (Nikon FN1) with 40X water-immersion objective. Fluorescence (mCherry) was detected with pE-300 and CCD camera (INFINITY 3S-1UR M/C). Patch pipettes (5-6 MΩ) were pulled from borosilicate glass capillaries (GBF-150-117-10; Sutter Instruments) with a micropipette horizontal puller (P-97, Sutter Instruments). Electrophysiological recordings were performed using a MultiClamp 700B amplifier (Molecular Devices) and acquired using a Digidata 1550B digitizer (Molecular Devices), sampled at 20 kHz for current clamp and 200 kHz for voltage-clamp, and filtered at 1 kHz. All data acquisitions were performed using pCLAMP 11.0.3 software (Molecular Devices). Stimulation of excitatory fibers was performed with a stimulating electrode (bipolar concentric electrode from Phymep and stimulator A365, World Precision Instruments). Intracellular solution based on K-gluconate (containing in mM: 128 KGlu, 20 NaCl, 1 MgCl2, 1 EGTA, 0.3 CaCl2, 2 Na2-ATP, 0.3 Na-GTP, 0.2 cAMP, 10 HEPES, pH 7.35) was used for recording basal electrophysiological properties (rest membrane potential (RMP), resistance, rheobase), spontaneous excitatory post-synaptic currents (sEPSC), paired-pulse ratio (PPR) and excitatory post-synaptic potential (EPSP)/spike coupling. Cesium-chloride intracellular solution (containing in mM: 150 CsCl, 2 MgCl2, 1 EGTA, 2 Na2-ATP, 0.2 Na-GTP, 0.2 cAMP, 10 HEPES, pH 7.35) was used to record spontaneous inhibitory post-synaptic currents (sIPSC). They were recorded in the presence of 10 mM CNQX (6-cyano-7-nitroquinoxaline-2,3-dione, Merck) and 50 mM D-AP5 (D-2-amino-5-phosphonovalerate, Tocris) to block AMPA and NMDA receptors, respectively. Recording of electrophysiological profile of DRD2-SPNs was performed in current clamp at resting membrane potential, with a series of 600 ms-duration depolarizing current steps starting at -150 pA with 10 pA increment. For sEPSC recordings, neurons were clamped at - 70mV. For sIPSC recordings, neurons were clamped at -60mV. For paired-pulse ratio, excitatory fibers were stimulated twice with an interval of 50 ms, while EPSC were monitored in voltage clamp. For the EPSP/Spike (E-S) coupling protocol, neurons were current clamped at -60 mV and stimulations of excitatory inputs were applied at 0.125 Hz with 6 different intensities (5 sweeps / intensity) chosen by the experimenter. EPSP slopes were measured individually off-line and the firing probability was plotted as a function of the EPSP slope (mV/ms). Data were analyzed offline using Clampfit 11.0.3 (Molecular Devices).

### Stereotaxic injections

Annk1^lox/lox^ animals were anaesthetized with isoflurane and received 10 mg.kg^-1^ intraperitoneal injection (i.p.) of Buprécare® (Buprenorphine 0.3 mg) diluted 1/100 in NaCl 9 g.L^-1^ and 10 mg.kg^-1^ of Ketofen® (Ketoprofen 100 mg) diluted 1/100 in NaCl 9 g.L^-1^, and placed on a stereotactic frame (Model 940, David Kopf Instruments, California). We bilaterally injected 0.6 (DS) or 0.3 (NAc) µl of virus (AAV9.CMV.HI.eGFP-Cre.WPRE.SV40 ou AAV5.CMV.HI.eGFP-Cre.WPRE.SV40 and GFP controls, (titer >10^13^ vg.mL^-1^, working dilution 1:10 into the DS (L = +/-1.75; AP = +0.6; V = -3.5, and -3 in mm) or the NAc (L=+/-1; AP=+1.55, V=-4.5) at a rate of 100 nl.min^-1^. The injection needle was carefully removed after 5 min waiting at the injection site and 2 min waiting half way to the top.

### Behavioral assays

#### Haloperidol-induced catalepsy

mice were injected with haloperidol (0.5 mg.kg^-1^, i.p.). Catalepsy was measured at several time points, 45-180 min after haloperidol injection. Animals were taken out of their home cage and placed in front of a 4-cm-elevated steel bar, with the forelegs upon the bar and hind legs remaining on the ground surface. The time during which animals remained still was measured. A behavioral threshold of 180 seconds was set so the animals remaining in the cataleptic position for this duration were put back in their cage until the next time point.

#### T-maze

Mice were tested for learning and cognitive flexibility in a gray T maze (arm 35-cm length, 25-cm height, 15-cm width). All mice were mildly food deprived (85-90 % of original weight) for 3 days prior to starting the experiment. The first day mice were placed in the maze for 15 min for habituation. Then, mice underwent 3 days of training with one arm reinforced with a highly palatable food pellet (HFHS, cat n. D12451). Each mouse was placed at a start point and allowed to explore the maze. It was then blocked for 20 seconds in the explored arm and then placed again in the starting arm. This process was repeated 10 times per day. At the end of the learning phase all mice showed a > 70 % preference for the reinforced arm. The average number of entries in each arm over 5 trials was plotted. Two days of reversal learning followed the training phase during which the reinforced arm was changed and the mice were subjected to 10 trials per day with the reward in the arm opposite to the previously baited one.

#### Operant conditioning experiments

The operant conditioning experiments were carried out in mouse operant chamber (Phenomaster, TSE Systems GmbH, Bad Homburg, Germany). Each chamber was composed by 2 lever-press, located 3 cm lateral to the food cup, one randomly selected as active and the other as inactive, one house light, a food dispenser and a food magazine between the 2 levers. The beginning of the session was concomitant with the turning on of the house light for 3 seconds, and a pellet delivery. The session ended after 60 minutes. Seven days before the beginning of the experiment mice were transferred in behavioral room and maintained in an environment with controlled temperature and humidity. Five days before the start of conditioning all the mice were food-restricted to maintain their weight at 90% of the beginning experiment original weight. The food restriction was maintained until the progressive ratio to facilitate the acquisition of the task. For NAc groups, mice where switched *ad libitum* food for a second progressive ratio. During the operant conditioning sessions animals were presented with 20 mg dustless precision highly palatable pellets (peanut butter flavored sucrose tablet, 5UTL Test Diet). The training session started with a fixed ratio of 1 (FR1) reinforcing schedule. During this period, animals had press one time on the active lever press to receive a pellet. Pressing on the active lever press resulted in the delivery of one pellet concomitant with a 2-second poke-light. Each poke in the active hole was followed by a 10-second time-out period during which the pokes were inactive, independently of the reinforcing schedule. The FR1 was followed by up to 10 days of FR5, during which mice were fasted at 90% of their original weight. When controls grouped reached the criterion, which was maintaining at least 70% responding in the active hole, a minimum of 10 reinforces per session in three consecutive days. Ankk1^lox/lox^, Ankk1^Δ-DR2 N^, Ankk1^GFP-NAc^, and Ankk1^Δ-NAc^ and were also subjected to a progressive ratio (PR) schedule during which the response requirements increased systematically within the session, after each reinforce and time out. The PR schedule lasted for 1 h and respected the following series: 1, 2, 3, 4, 5, 6, 7, 8, 10, 12, 14, 16, 18, 20, 22, 24, 28, 32, 36, 40, 44, 48, 52, 56, 64, 72, 80, 88, 96, 104, 112, 120, 128, 136, 144, 152, 160, 168, 176, 184, 192, 200, 208, 216, 224, 232, 240, 248, 256, 264, 272, 280, 288, 296, 304, 312, 320, 328, 336, 344, 352, 360, 368, 376, 384, 392, 400, 408, 416, 424, 432, 440, 448, 456, 464.

### Indirect calorimetry analysis

All mice were monitored for metabolic efficiency (Labmaster, TSE Systems GmbH, Bad Homburg, Germany). After an initial period of acclimation in the calorimetry cages of at least two days, food and water intake, whole energy expenditure (EE), oxygen consumption and carbon dioxide production, respiratory quotient (RQ=VCO2/VO2, where V is volume) and locomotor activity were recorded as previously described^83^. Additionally, fatty acid oxidation was calculated as previously reported^83^. Reported data are the results of the average of the last three days of recording. Before and after indirect calorimetry assessment, body mass composition was analyzed using an Echo Medical systems’ EchoMRI (Whole Body Composition Analyzers, EchoMRI, Houston, USA).

### Statistical analyses

Compiled data are always reported and represented as mean ± SEM., with single data points plotted (single cell our mouse). Data were statistically analyzed with GraphPad Prism 9. Normal distribution was tested with Anderson-Darling, D’agostino Pearson test, Shapiro-Wilk test and Kolmorogrov-Smirnov. Data were analyzed with Two-tailed Mann-Whitney, unpaired Student’s T-Test, one-way ANOVA, two-way ANOVA or repeated-measures ANOVA, as applicable and Holm-Sidak’s post-hoc test for two by two comparisons. All tests were two-tailed. Significance was considered as p<0.05. Detailed statistical results are reported in **Supplementary Table 3**.

### Human participants

A total of thirty-six subjects were recruited from the greater New Haven, Connecticut area via flyers or social media advertisements. Subjects were enrolled in the pilot study based on BMI (<26) and underwent indirect calorimetry measurement and genotyping in different studies. All subjects provided written informed consent at the first visit and the study was approved by the Yale Human Investigation Committee.

#### Indirect calorimetry analysis

Participants were instructed to fast for at least 4 hours; abstain from alcohol, caffeine, and moderate exercise for at least 2 hours; and abstain from vigorous exercise for at least 14 hours prior to the session. Resting Respiratory Quotient (RQ) was measured for 45-min by a ParvoMedics TrueOne metabolic cart (ParvoMedics, Salt Lake City, UT) following recommended best practices (Compher et al., 2006). All indirect calorimetry measurements were first prepared for analysis by discarding the first 5-min to allow for proper equipment calibration. To determine steady state energy expenditure, the measurement was assessed with 5-min rolling windows to identify the time frame with the lowest coefficient of variance of both VCO_2_ and VO_2_ for each subject (Compher et al., 2006). The average was then taken of this 5-min time frame and used as the steady state measure. Before and after the measurement, participants were asked to rate their internal state including hunger, fullness, thirst, and others. Two subjects were excluded from the analysis: one did not complete the 45-min session due to technical issues, the other fell asleep during the assessment.

#### Genotyping

Saliva samples were obtained from subjects using Oragene-Discover collection kit (OGR-600) and DNA extraction was performed as instructed by the manufacturer (DNA Genotek, Canada). Genotyping was performed at Transnetyx (Cordova, TN) using a qPCR-based system to detect the presence or absence of a target sequence within each sample. The HuAnkk1-1 MUT probe was used and was targeted to the rs1800497 SNP variant found within the genomic human Ankk1 sequence. The HuAnkk1-1 MUT probe is targeted to differentiate between the G>A SNP variant in the human Ankk1 gene and detects the presence of either the wild-type or mutant SNP in the human cassette that is randomly integrated into the mouse genome. This assay utilizes a multiplex design strategy where two fluorescently labeled reporter oligos compete for the binding site and have a common forward and reverse primer. Seventeen were Taq1A A1 allele carriers (A1+) and 19 were non-carriers (A1)-. Of the 17 A1+, only one were homozygotes, so we collapsed across heterozygotes and homozygotes.

#### Statistical analysis

GraphPad Prism 9 was used to visualize data and compare the baseline characteristics using Independent-samples t-tests. Anomalies were defined by 2* standard deviation and two such outliers were excluded, resulting in total 32 subjects included in the results. Linear regression analysis was conducted to detect the difference in RQ between subjects with A1+ and A1-while controlling for age, sex, BMI, and study using Python 3.8.5 (statsmodels package).

