## Supplemental figures and tables for "The addiction-susceptibility TaqIA/Ankyrin repeat and kinase domain containing 1 kinase (ANKK1) controls reward and metabolism through dopamine receptor type 2 (DR2)-expressing neurons"

### Supplementary Material

**Enrica Montalban *et. al***

**List of Figures**

**Supplementary figures**

**Supplementary figure 1.** Targeting construct Ankk1 floxed

**Supplementary figure 2.** Consequences of Ankk1 downregulation in DR2 neurons on operant conditioning learning and motivation

**Supplementary figure 3**. Consequences of Annk1 downregulation in DS and NAc on *Ankk1* and *Dr2* m-RNA expression and cataleptic response to haloperidol

**Supplementary figure 4**. Consequences of Ankk1 downregulation in the NAc on impulsive behavior, motivation and binge eating.

**Supplementary figure 5**. Consequences of Ankk1 downregulation in DS on striatal behavior and energy metabolism

**Supplemental Table 1:** Basic membrane properties of DRD2-SPNs neurons in Drd2-CRE and Ankk1$^{\Delta\text{-}DR2N}$ mice.

**Supplementary Table 2.** Participant Characteristics

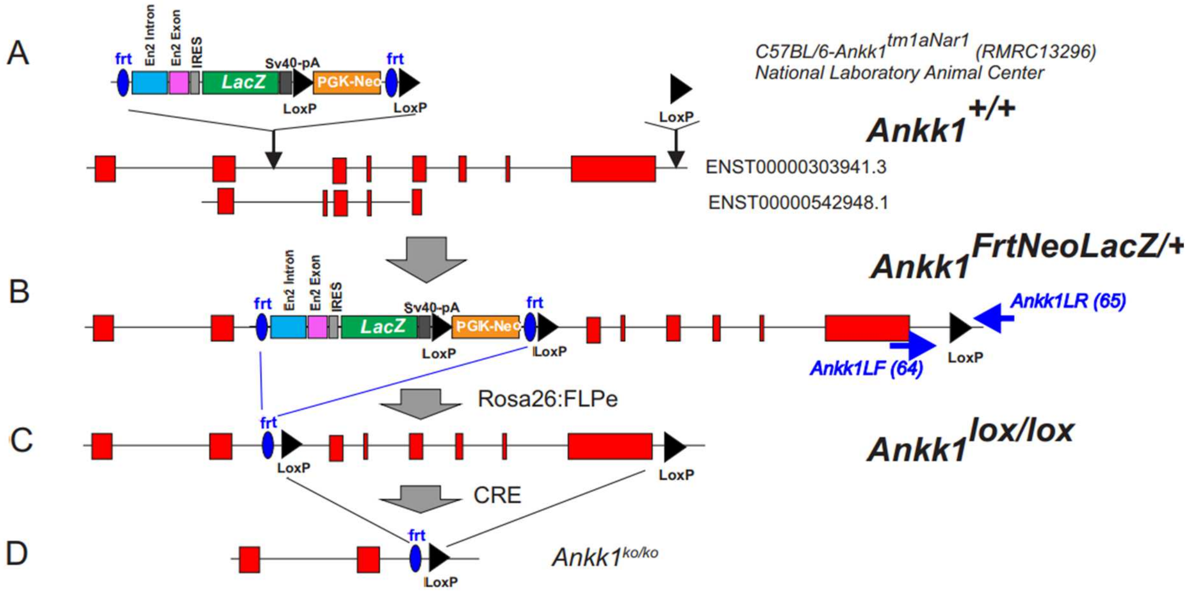


**Supplementary figure 1. Targeting construct of Ankk1 floxed line**. **A.** C57BL/6-Ankk1$^{tm1aNarl}$ were obtained by the insertion of a FRT-*En2* Intron-*En2* Exon-IRES-LacZ-***LoxP***-PGK-Neo FRT:***LoxP*** between Exon 2 and 3, in addition to an insertion of a LoxP site after exon 8. **B.C** Ankk1$^{FrtNeoLacZ/+}$ were bred with B6N.129S4-Gt(ROSA)26Sor*tm1(FLP1)Dym/J (Rosa26::FLPe Knock in)* expressing the flippase under the ubiquitous Rosa26 promoter. **D**. The resulting mice Ankk1$^{lox/lox}$ mice contains a FRT-LoxP site between Exon 2 and 3 in addition to an insertion of a LoxP site after exon 8.

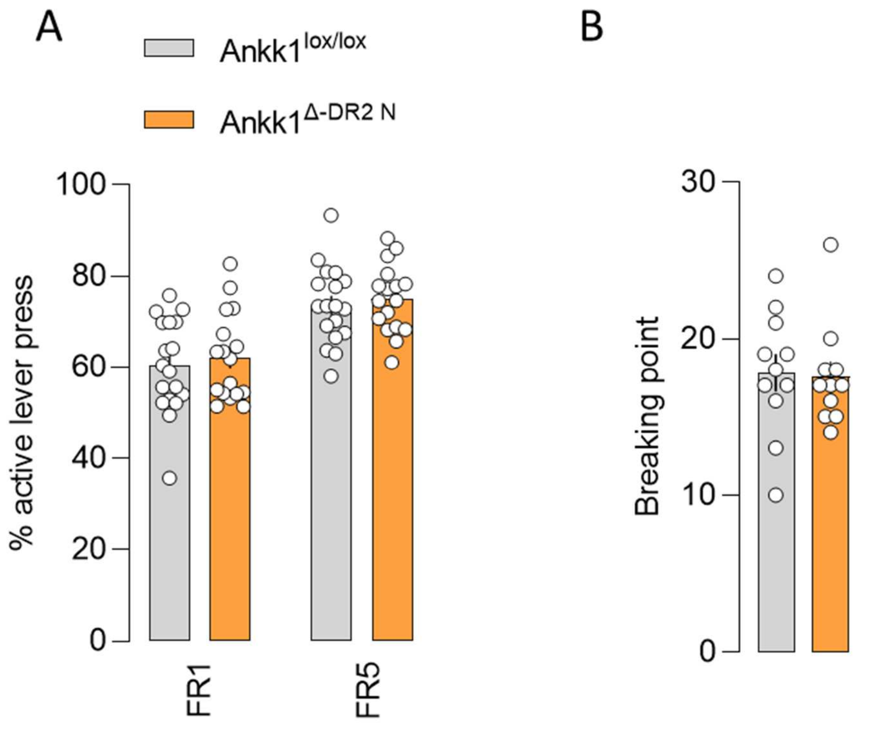


**Supplementary figure 2. Consequences of Ankk1 downregulation in D2 neurons on operant conditioning learning and motivation:** Ankk1$^{lox/lox}$ and Ankk1$^{\Delta\text{-DR2 N}}$ show comparable average of % active lever press per session during each period of conditioning (**A)** and motivation as showed by the breaking point in **(B).** Data are expressed as mean ± SEM. n=18 for conditioning and n=11 for breaking point.

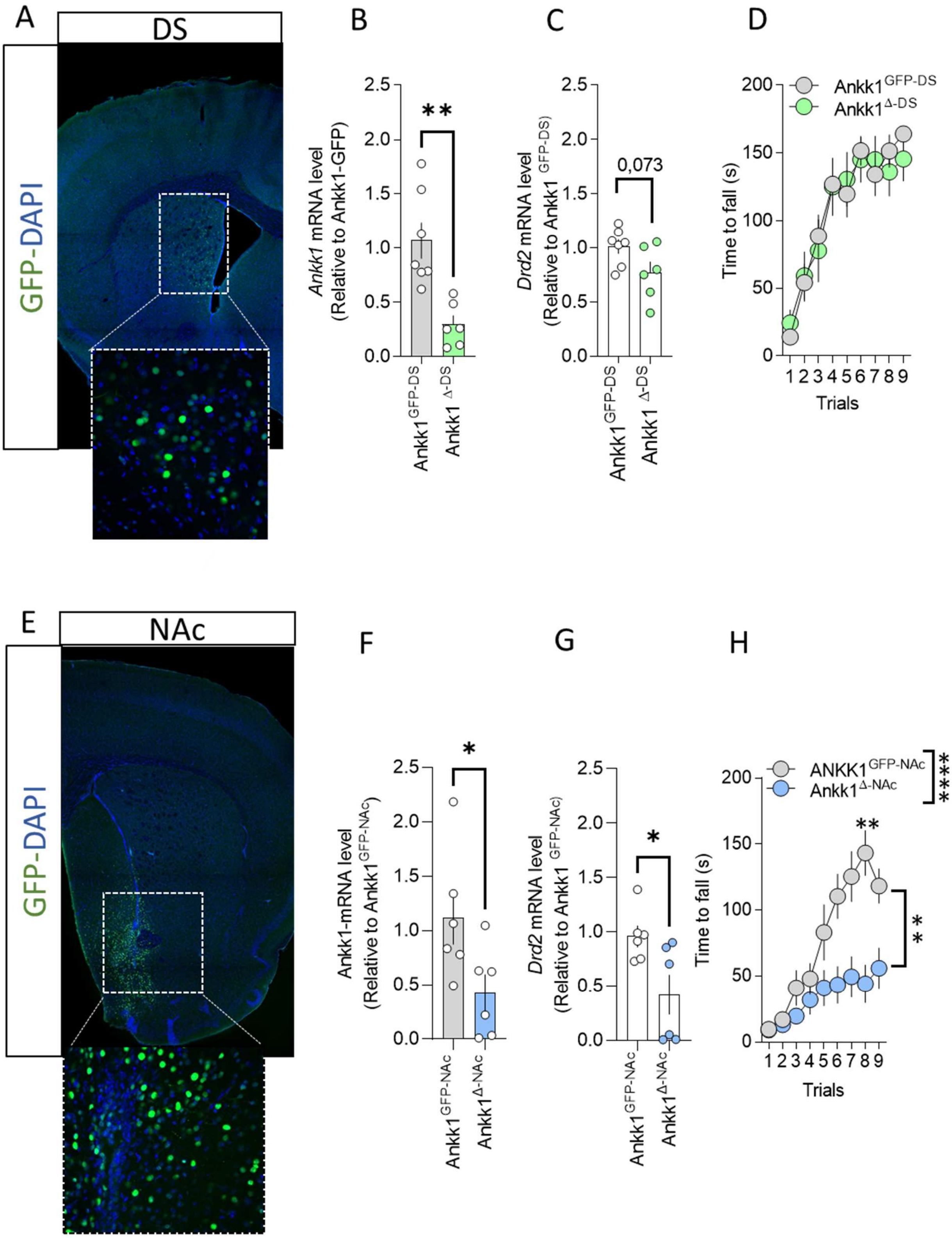


**Supplementary figure 3**. **Consequences of Annk1 downregulation in DS and NAc on Ankk1 and *Dr2* m-RNA expression, and cataleptic response to haloperidol**. **A-E** Representative histological controls for AAV-CRE expression. Mosaic of confocal images in representative mice used for viral injection DAPI (blue)/AAV-CRE (green). Ankk1 lox/lox mice were stereotactically injected in the DS (**A**) or the NAc (**E**). Scale bar, 200 µm. Inset: higher magnification of a different section, scale bar 150 µm. **B-E** Real time control for AAV-CRE dependent downregulation of *Ankk1* m-RNA in DS (**B**) and NAc (**F**). The mRNA was purified from NAc and DS of Ankk1$^{GFP-}$ and Ankk1$^{\Delta-}$ mice, and analyzed by qRT-PCR. The expression levels were calculated by the comparative ddCt method with RPL19 as an internal control. Data points are individual results from different mice (n = 6-7 per group). Means ± SEM are indicated. Statistical analyses are performed with two-tailed Mann-Whitney's test, p = 0.0012 for DS and p=0.0260 for NAc. **C-G** *Dr2* m-RNA level Ankk1$^{GFP-}$ mice and Ankk1$^{\Delta-}$ mice in DS (**C**) and NAc (**G**). Data points are individual results from different mice (n = 6-7 per group). Means ± SEM are indicated, Statistical analyses are performed with two-tailed Mann-Whitney's test, p = 0.073 for DS and p=0.0260 for NAc. **D-H** The effects of *Ankk1* and *Dr2* m-RNA downregulation on DR2 function were investigated by evaluating the immobility 45-180 min after haloperidol injection (0.1 mg.kg$^{-1}$, i.p.) for DS (**D**) and NAc (**H**). n=10-7 for DS and 9-13 for NAc Satistics NAc: Two-way ANOVA, Interaction p<0.0001, time p<0.0001 genotype p<0.0001, Sidak's post-hoc test, line 6 p=0.0033, line 7 p=0.0005 Line 8 p<0.0001, line 9 p=0.0081

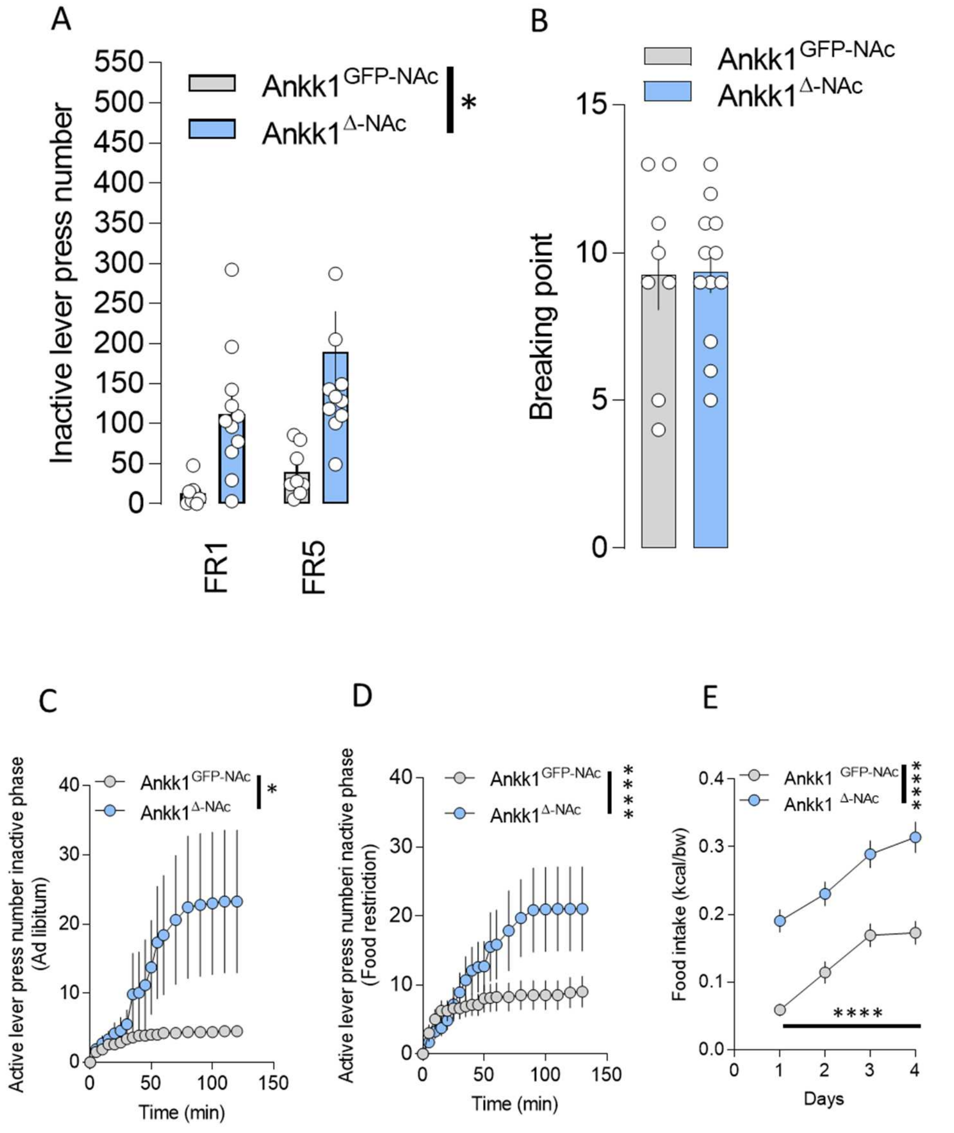


**Supplementary figure 4**. **Consequences of Ankk1 downregulation in the NAc on impulsive behavior, motivation and binge eating**. **A** Ankk1 $^{\Delta\text{-NAc}}$ showed a significantly higher number of inactive lever press than Ankk1$^{\text{GFP-NAc}}$ mice. Histogram shows the average inactive lever press per session of operant conditioning. Data are expressed as mean ± SEM. n=8-11. Statistics: Interaction p=0.2589, Learning p=0.0293 Genotype p=0.0088. However, Ankk1$^{\text{GFP-NAc}}$ and Ankk1 $^{\Delta\text{-NAc}}$ mice show similar breaking point (**B**) in a progressive ratio paradigm. Data are expressed as mean ± SEM. n=8-12 **C** Ankk1 $^{\Delta\text{-NAc}}$ mice performed significantly more lever press during the inactive phase of the progressive ratio paradigm in both ad libitum (**C**) and fasted (**D**) conditions Statistics **C** Interaction p=0.0104, Time p=0.0844, Genotype p=0.2062. n=8-13. Statistics **D** Interaction p<0.0001, Time p=0.0042, Genotype p=0.2449. n=8-12. **E** Caloric intake of Ankk1$^{\text{GFP-NAc}}$ and Ankk1 $^{\Delta\text{-NAc}}$ mice during binge eating. Data are expressed as mean ± SEM, N=10-12. Interaction p<0.0001, time p<0.0001 genotype p<0.0001 Sidak's post-hoc test p<0.0001

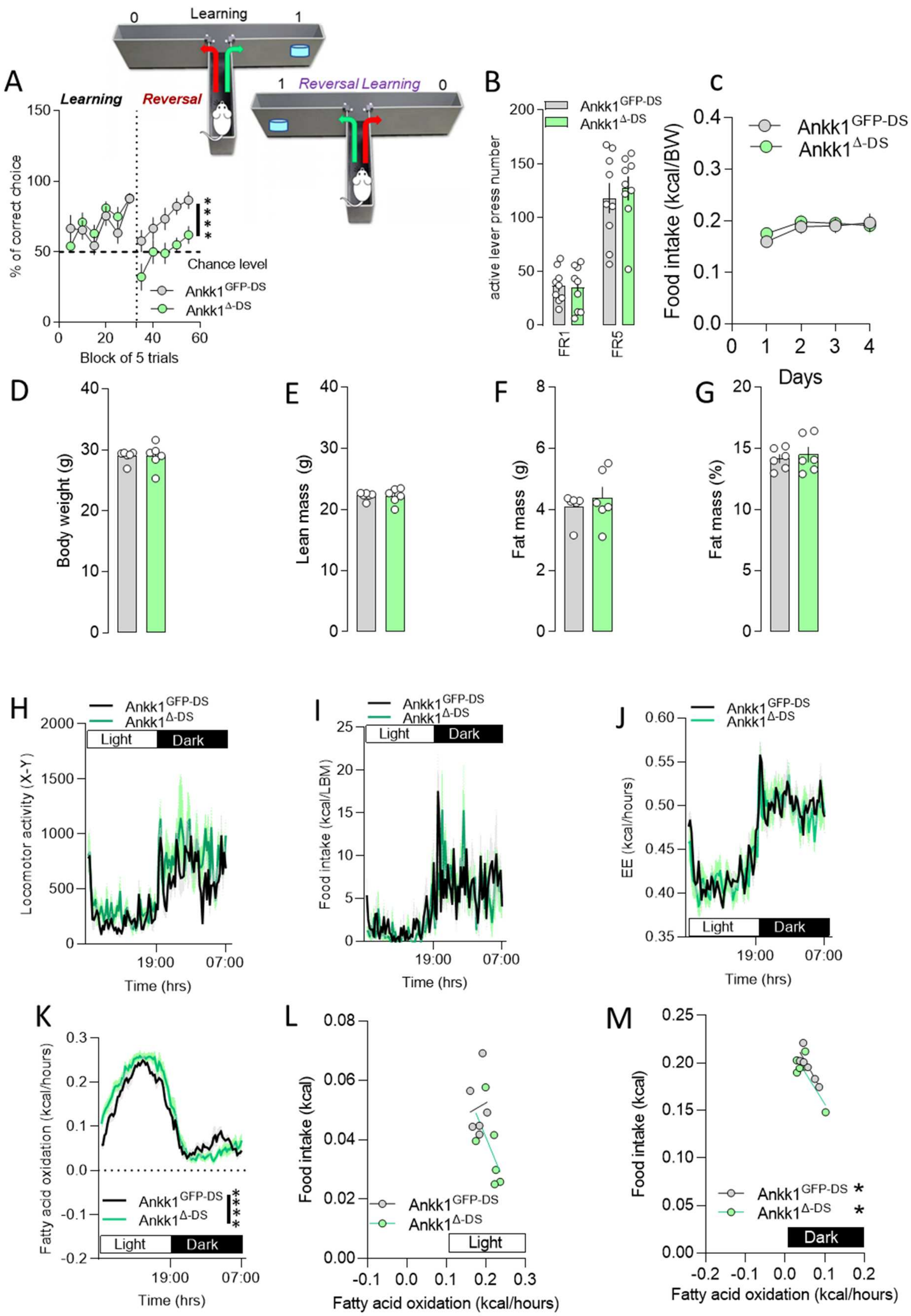

A
Learning
Reversal
Learning
Reversal Learning
% of correct choice
Chance level
Block of 5 trials
Ankk1GFP-DS
Ankk1Δ-DS
B
active lever press number
FR1
FR5
C
Food intake (kcal/BW)
Days
D
Body weight (g)
E
Lean mass (g)
F
Fat mass (g)
G
Fat mass (%)
H
Locomotor activity (X-Y)
Light
Dark
Time (hrs)
19:00
07:00
I
Food intake (kcal/LBM)
J
EE (kcal/hours)
K
Fatty acid oxidation (kcal/hours)
L
Food intake (kcal)
Fatty acid oxidation (kcal/hours)
M

**Supplementary figure 5**. **Consequences of Ankk1 downregulation in DS on striatal behavior and energy metabolism A**- Ankk1$^{GFP\text{-}DS}$ and Ankk1 $^{\Delta\text{-}DS}$ mice showed similar performances in the learning phase of a T-MAZE paradigm, however, Ankk1 $^{\Delta\text{-}DS}$ mice showed impaired performances during the reversal learning. Data are expressed as mean ± SEM. n=9-10. Statistics: Interaction p=0.4134, Learning p=0.0293, Genotype p=0.0088. **B** Average of active lever press per session during each period of conditioning, is comparable between Ankk1$^{GFP\text{-}DS}$ and Ankk1$^{\Delta\text{-}DS}$ n=9. Ankk1$^{GFP\text{-}DS}$ and Ankk1$^{\Delta\text{-}DS}$ also show comparable body weight (**D**), lean mass (**E**), fat mass (**F**), fat mass % (**G**), locomotor activity (**H**), caloric intake (**I**) and energy expenditure (**J**). Fatty acid oxidation is significantly different between Ankk1$^{GFP\text{-}DS}$ and Ankk1$^{\Delta\text{-}DS}$, Interaction p<0.0001, Time p<0.0001, Genotype p=0.2968, and while it does not correlate with food intake during the diurnal phase it does for the 2 groups during the night phase. Ankk1$^{GFP\text{-}DS}$ p=0.0296 and Ankk1$^{\Delta\text{-}DS}$ p=0.0453

**Supplemental Table 1: Basic membrane properties of DRD2-SPNs neurons in Drd2-CRE and Ankk1$^{\Delta\text{-DR2N}}$ mice.**

| | Drd2-CRE (n=11 neurons / 8 mice) | Ankk1$^{\Delta\text{-DR2 N}}$ (n=9 neurons / 7 mice) | p value (Mann-Whitney tests) |
|---|---|---|---|
| Rest membrane potential (mV) | -74.1 ± 2.6 | -76.3 ± 1.7 | p=0.5637 |
| Resistance (MΩ) | 208 ± 35 | 318 ± 58 | p=0.2014 |
| Rheobase (pA) | 101 ± 11 | 71 ± 10 | p=0.0753 |

**Supplementary Table 2. Participant Characteristics**

| Variables | A1- (N=19) | A1+ (N=13) | P value |
|---|---|---|---|
| Age (Years) | | | 0.7677 |
| N | 19 | 13 | |
| Mean ± SD | 27.8 ± 1.0 | 26.9 ± 5.8 | |
| Median (IQR) | 24 (4.5) | 26 (11) | |
| Sex, n (%) | | | 0.0014 |
| Female | 17 (89.5) | 5 (38.5) | |
| Male | 2 (10.5) | 8 (61.5) | |
| Race, n (%) | | | 0.0641 |
| White | 12 (63.2) | 4 (30.8) | |
| Black or African American | 1 (5.3) | 0 (0) | |
| Asian | 5 (26.3) | 8 (61.5) | |
| Others | 1 (5.3) | 1 (7.7) | |
| BMI (kg/m2) | | | 0.7273 |
| Mean ± SD | 22.3 ± 2.4 | 22.6 ± 1.9 | |
| Hours of sleep | | | 0.8896 |
| Mean ± SD | 7.1 ± 1.4 | 7.1 ± 1.1 | |
| Hours since last meal | | | 0.5254 |
| Mean ± SD | 11.8 ± 3.1 | 11.0 ± 3.6 | |
| Hip circumference (cm) | | | 0.7288 |
| Mean ± SD | 97.5 ± 6 | 96.8 ± 4.2 | |
| Waist circumference (cm) | | | 0.2525 |
| Mean ± SD | 75.1 ± 7.2 | 78.3 ± 8 | |
| Waist/hip ratio | | | 0.0879 |
| Mean ± SD | 0.8 ± 0.1 | 0.8 ± 0.1 | |
| FM (kg) | | | 0.3017 |
| Mean ± SD | 17 ± 6 | 14.9 ± 4.3 | |
| FMI (kg/m2) | | | 0.0515 |
| Mean ± SD | 6.7 ± 2.5 | 5.1 ± 1.5 | |
| VAT (l) | | | 0.6055 |
| Mean ± SD | 1.3 ± 1.7 | 1.5 ± 1.2 | |

Independent t-tests were performed to compare A1+ and A1- groups. Abbreviations: SD: standard deviation, IQR: interquartile range, BMI: body mass index, FM: fat mass, FMI: fat mass index, VAT: visceral adipose tissue.
